# Dynamic changes in the regulatory T cell heterogeneity and function by murine IL-2 mutein

**DOI:** 10.1101/669978

**Authors:** Daniel R. Lu, Hao Wu, Ian Driver, Sarah Ingersoll, Sue Sohn, Songli Wang, Chi-Ming Li, Hyewon Phee

## Abstract

The therapeutic expansion of Foxp3^+^ regulatory T cells (Tregs) shows promise for treating autoimmune and inflammatory disorders. Yet, how this treatment affects the heterogeneity and function of Tregs is not clear. Using single-cell RNA-seq analysis, we characterized 31,908 Tregs from the mice treated with a half-life extended mutant form of murine IL-2 (IL-2 mutein, IL-2M) that preferentially expanded Tregs, or mouse IgG Fc as a control. Cell clustering analysis revealed that IL-2M specifically expands multiple sub-states of Tregs with distinct expression profiles. TCR-profiling with single-cell analysis uncovered Treg migration across tissues and transcriptional changes between clonally related Tregs following IL-2M treatment. Finally, we identified IL-2M-expanded Tnfrsf9^+^Il1rl1^+^ Tregs with superior suppressive function, highlighting the potential of IL-2M to expand highly suppressive Foxp3^+^ Tregs.

**One Sentence Summary:** Single-cell analysis revealed that IL-2 mutein treatment expanded multiple sub-states of Tregs with a highly suppressive function in mice.

## Introduction

Foxp3^+^ regulatory T cells (Tregs) play a fundamental role in immunosuppression and immune tolerance, and there is great interest in harnessing Treg populations to treat autoimmune and inflammatory disorders. The differential expression of transcription factors, costimulatory receptors, chemokine receptors, and secreted effectors in quiescent and activated Tregs suggests that the heterogeneous Treg states exist and perform distinct functions(*1–3*). Moreover, non-lymphoid tissue Tregs acquire unique phenotypes different from lymphoid-tissue Tregs, suggesting that the anatomical location of Tregs contributes to their heterogeneity(*4, 5*).

Recently, low-dose Interleukin-2 (IL-2) therapies have been tested to induce tolerance in patients with autoimmunity and inflammatory disorders(*6–11*). Although the low-dose IL-2 therapies expand Tregs, their effect has been limited by concomitant increases in conventional effector T cells and natural killer cells. To improve selectivity and pharmacokinetics of low-dose IL-2, alternative modalities have been considered(*12*). However, it is not clear how IL-2-based therapies impact Treg heterogeneity in diverse tissues. Since the goal of Treg-targeted therapies is to expand Treg-mediated tolerance at the proper anatomical locations, it is critical to understand how IL-2-mediated expansion impacts the phenotypic and functional heterogeneity of Tregs in lymphoid and non-lymphoid tissues.

Thymic-derived Foxp3+ Tregs undergo T cell receptor (TCR) dependent antigen priming and activation-induced expansion in lymphoid organs followed by extravasation into peripheral tissues, where they acquire tissue-specific tolerogenic phenotypes. Given the complex migratory patterns of Tregs, it is unclear how IL-2-mediated therapy affects Tregs within and across tissues. TCR-sequencing combined with single-cell profiling provides an opportunity to measure IL-2-induced Treg differentiation and movement by tracing the transcriptional conversions and trafficking patterns of clonal lineages.

To better understand the impact of the IL-2-mediated Treg expansion therapy on Foxp3^+^Treg heterogeneity in lymphoid and non-lymphoid tissues, we profiled mouse spleen, lung and gut Tregs using single-cell RNA-seq (scRNA-seq) with TCR-sequencing under murine IL-2 mutein (IL-2M) stimulation or homeostatic (mouse IgG Fc isotype control-treated) conditions. Comparison of resting, primed/activated, and activated Treg states from different tissues revealed unique gene signatures shared between spleen and lung Tregs, as well as distinct activation profiles of gut Tregs. Administration of murine IL-2M dramatically changed the landscape of Tregs in the spleen and the lung, while maintaining tissue-specific identity in the gut. TCR-profiling coupled with scRNA-seq revealed gene expression dynamics governing Treg differentiation after IL-2M treatment and uncovered a migratory axis across tissues. Additionally, we identified a population of activated *Tnfrsf9^+^Il1rl1*^+^ Tregs in mice that expands following IL-2M and suppresses convention T cells robustly *in vitro*. Overall, our experiments provide new insights into the relationships between Foxp3^+^ Treg activation states and their phenotypic heterogeneity in different tissues during homeostasis and after murine IL-2M stimulation.

## Results

### A half-life extended mutant form of murine IL2 expands CD25^+^Foxp3^+^ Tregs in vivo

To determine the specific role of mouse IL-2 in Foxp3^+^ Tregs in mice, a half-life extended mutant form of murine IL-2 (IL-2 mutein, IL-2M) was generated as an Fc fusion protein. Administration of murine IL-2M to C57BL/6 mice with different doses revealed that 0.33 mg/kg of murine IL-2M specifically increased Foxp3^+^ Tregs in the spleen, lymph nodes, blood and lung, while it did not affect CD25^−^Foxp3^−^ T conventional cells (Tconv) (Fig. 1A, Fig. S1A-D). Murine IL-2M increased percentages and cell numbers of CD25^+^Foxp3^−^ activated CD4 T cells slightly at a higher dose (Fig. S1A), likely due to the presence of high affinity IL2R (IL2Rαβγ in CD25 expressing cells(*13, 14*). Expansion of CD25^+^Foxp3^+^ Tregs by IL-2M was comparable to IL-2/anti-IL2 antibody (JES6-1) conjugate (IL-2C) which was previously reported to expand Tregs(*15*) (Fig. S2C), but IL-2C increased more CD25^+^Foxp3^−^ activated CD4 T cells compared with IL-2M (Fig. S2D).

**Fig. 1.**
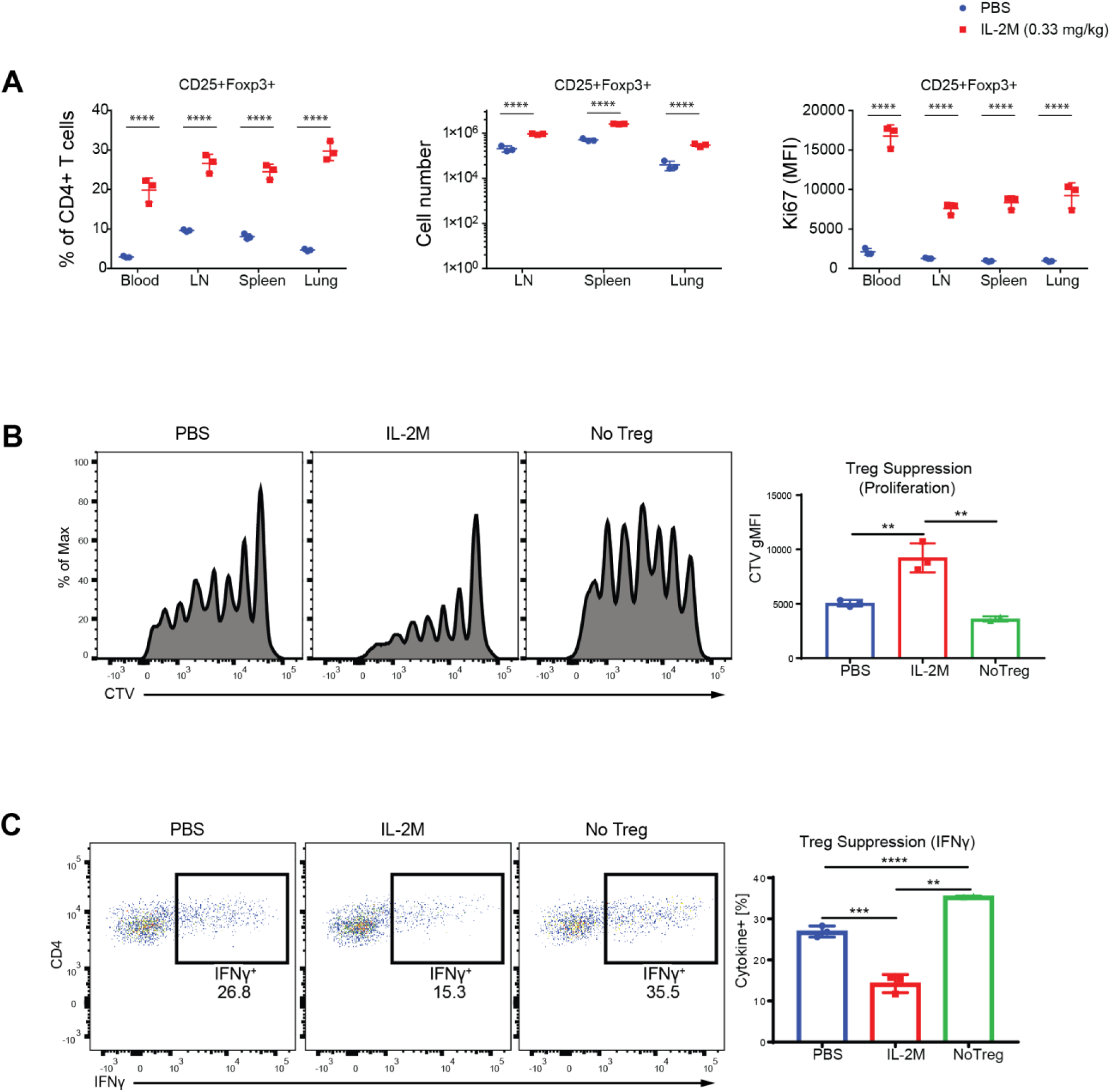
A half-life extended mutant form of the murine IL-2M expands CD25^+^Foxp3^+^ Tregs *in vivo* and increases their suppressive activity. (**A**) Effects of IL-2M (0.33 mg/kg) on the percentages of CD25^+^Foxp3^+^ Tregs within CD4^+^ T cells (*left panel*) and cell numbers of CD4^+^CD25^+^Foxp3^+^ Tregs (*middle panel*) from the blood, spleen, lymph nodes and lung. Effects of IL-2M (0.33 mg/kg) on the proliferation of CD4^+^CD25^+^Foxp3^+^ Tregs assessed by expression of Ki67 (mean fluorescence intensity) (*Right panel*). Effects of IL-2M on other compartments of CD4 T cells are shown in Fig. S1 A to D. Results are representative of at least three independent experiments. Experiments were performed using 3 mice per each group. *Statistics*; Two-way ANOVA for multiple comparisons. *0.01< *p* <0.05, **0.001< *p* <0.01, ***0.0001< *p* <0.001, \*\*\*\**p* < 0.00001. (**B**) *In vitro* suppression assays using sorted Foxp3-EGFP+ Tregs from the spleen of either PBS or murine IL-2M-treated Foxp3EGFP mice. Naïve T cells were MACS sorted using Miltenyi naïve T cell isolation kit, labeled with cell tracer violet (CTV), then mixed with irradiated APC in the absence (No Tregs) or presence of Tregs (PBS or IL-2M). Tregs were cocultured then with naïve T cells and APCs under Th1 skewing condition. Shown are representative images of proliferation of Tconv indicated as dilution of CTV. (**C**) Suppression of IFNγ generation from Th1 effector cells by Tregs. *In vitro* Treg suppression assay was performed as described in **B**, and expression of IFNγ from Tconv was measured by intracellular staining. Results are representative of two independent experiments. *Statistics*; One-way ANOVA for multiple comparisons. **0.001< *p* <0.01, ***0.0001< *p* <0.001, \*\*\*\**p* < 0.00001.

*In vivo* expanded Tregs by IL-2M displayed superiority in suppressing the proliferation of Th1 effector cells (Fig. 1B) as well as generation of IFNγ from Th1 effector cells *in vitro* compared with Foxp3^+^ Tregs isolated from spleens of PBS-treated mice (Fig. 1C). While these results revealed that highly suppressive Tregs expanded after IL-2M treatment *in vivo*, it was not clear how IL-2M impacted on heterogeneity of Tregs towards functional suppression in lymphoid and non-lymphoid tissues. We therefore sought to profile the molecular phenotypes expressed by Tregs to understand how IL-2M elevates Treg expansion and suppression in mice.

### Isolation and scRNA-seq of Tregs from spleen, lung and gut

The therapeutic success of Treg expansion depends on the efficacy of Treg immunosuppression at specific anatomical sites. To characterize the heterogeneity of Tregs and assess how IL-2M reshapes this diversity, we performed scRNA-seq on Foxp3^+^ Tregs from multiple tissues at steady-state and after IL-2M treatment. CD4^+^eGFP^+^ single cells were sorted from the spleen, lung, and gut of Foxp3-eGFP mice and single cell libraries were prepared using the 10x Chromium platform (Fig. 2A). After filtering and cross-sample normalization using Seurat(*16*), we recovered 17,097 spleen Tregs, 10,353 lung Tregs, and 4,458 gut Tregs across three replicates with roughly equivalent Tregs in mouse IgG Fc isotype control (Iso)- and IL-2M-treated conditions (16,152 and 15,756 cells, respectively). Single-cell molecule and gene recovery rates showed good concordance between Iso- and IL-2M-treated cells (Fig. S3A-B). To establish that we had accurately sorted and sequenced Tregs, we additionally performed scRNA-seq on CD4 conventional T cells sorted using FACS. Comparison of Tregs and Tconvs confirmed that Tregs and Tconvs expressed distinct transcriptional profiles and clustered into distinct groups (Fig. S4A). Additionally, Tregs expressed higher transcript levels of established Treg genes such as *Foxp3*, *Il2ra*, *Ctla4*, *Ikzf2*, and *Nrp1*, while both cell types expressed similar levels of *Cd4* (Fig. S4, B and C).

**Fig. 2.**
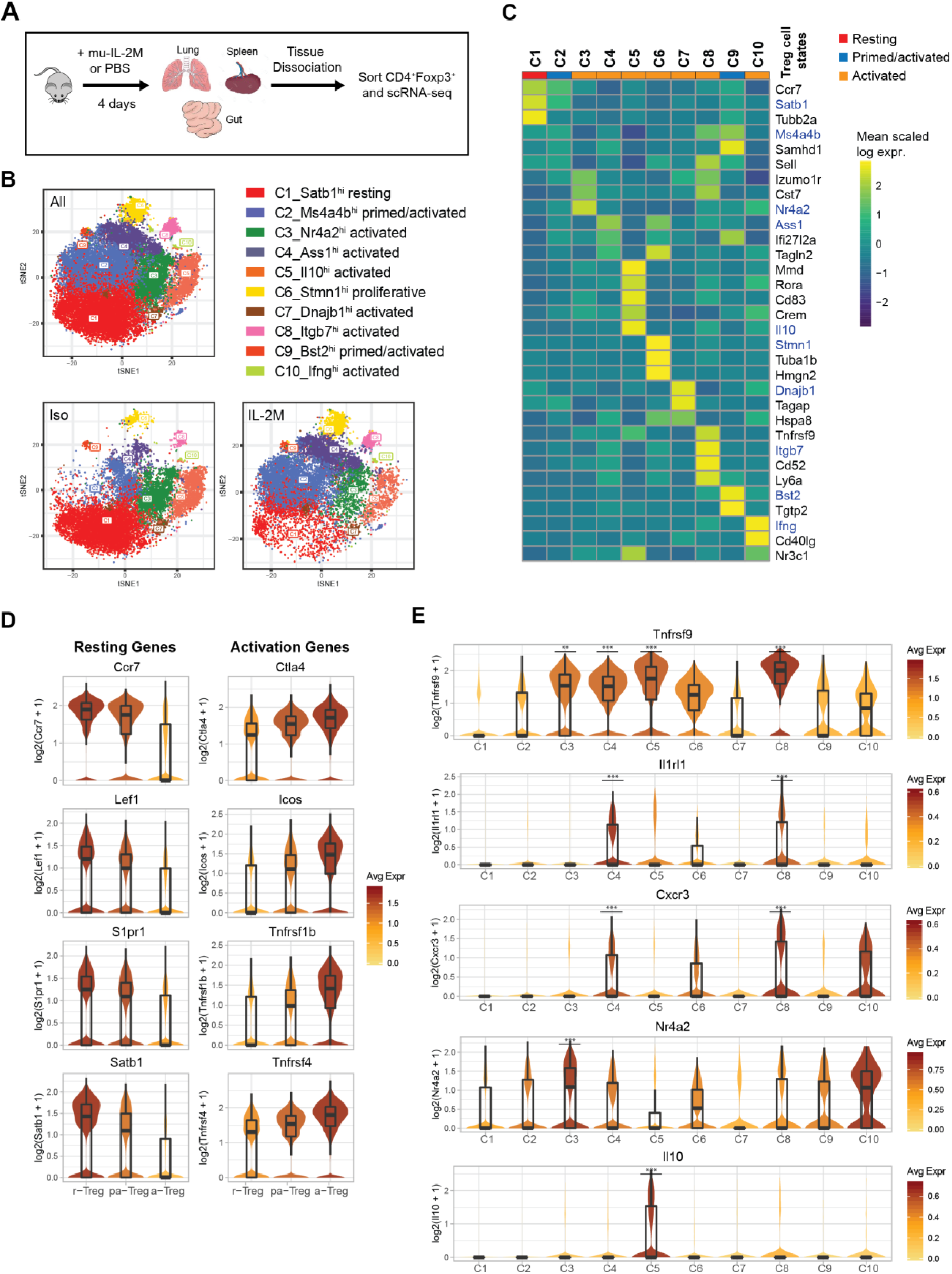
Classification of heterogeneous Treg resting, primed, and activation states across tissues. **(A)** Experimental workflow for interrogation of Tregs in the spleen, lung, and gut following administration of IL-2M or mouse IgG Fc control (Iso) by scRNA-seq. **(B)** t-SNE projection of all Tregs (top), Tregs from Iso-treated mice (bottom left), or Tregs from IL-2M-treated mice (bottom right) using K-nearest-neighbor graph-based clustering. Individual cells are colored by cell state cluster assignments. **(C)** Heatmap depicting the mean expression (Z-score) of most representative signature genes (rows) for each cluster (columns). Genes shown in blue text with an asterisk are genes used to characterize Treg states. **(D)** Violin plots depicting gradual downregulation of classical resting Treg markers (left) and gradual upregulation of classical activated Treg markers (right) in progressively higher Treg activation states. **(E)** Violin plots showing marker genes associated with sub-populations of activated Tregs. *Statistics*; \*\**p*<0.01, \*\*\**p*<0.001, Wilcoxon rank sum test.

### Molecular characterization of resting, primed, and activated Treg cell states across all tissues

Previous scRNA-seq profiling efforts have identified resting, early-activated (primed/activated), and terminal effector states along the Treg cell differentiation continuum(*4, 17*),(*18–20*). We sought to organize all Tregs from our dataset into different cellular states to generate a single-cell Treg classification schema, which could then define cellular variation across tissues and after IL-2M stimulation. To do this, we first impartially defined cell states using unsupervised clustering, then identified highly representative signature genes from each cell state to anchor our cell state assignments in the context of known Treg biology.

Cell clustering using the graph-based Louvain algorithm from Seurat(*21*) identified ten Treg states that were differentially distributed in isotype control (Iso)- and IL-2M-treated mice (Fig. 2B). These clusters were evenly represented in all replicates, indicating that they represent biological heterogeneity and not batch-specific processing artifacts (Fig. S5, A and B).

Signature genes were then defined for each cell state to understand transcriptional differences between Treg populations (see Methods) (Fig. 2C, Fig. S6A). Cell states identified by clustering could be grouped into resting, primed/activated, and activated Treg populations based on known lymphoid-associated and activated gene expression. Resting Tregs (C1) have high expression of lymphoid-tissue homing receptors (*Ccr7, S1pr1, Sell*) and Treg-establishing transcriptional regulators (*Lef1(22)*, *Satb1(23)* and *Klf2(24)*) and low activation marker expression (*Ctla4*, *Icos*, *Tnfrsf1b*, *Tnfrsf4*) (Fig. 2D, Fig. S7A), suggesting that they either have not yet undergone TCR-mediated activation or have a central memory-like phenotype.

Two primed/activated Treg states (C2 and C9), express intermediate levels of both resting and activation gene sets, suggesting they may be transitioning toward an activated phenotype (Fig. 2D, Fig. S7A). In support of this observation, primed/activated Tregs are distinguishable from C1 resting Tregs with higher expression of Treg-specific genes that stabilize inhibitory activity of Treg (C2: *Ikzf2(25)* and S100 family proteins) or inflammatory response mediators (C9: *Stat1*, *Cxcl10*, and interferon-response genes) (Fig. S7, B and C). Both C2 and C9 have high *Ms4a4b* expression (Fig. S7D), which modulates activation by regulating proliferation and augmenting co-stimulation through GITR(*26, 27*). Furthermore, C2- and C9-Tregs could be distinguished from each other, as C2-Tregs express more *Nrp1* (which is expressed highly in natural Tregs(*28*) and promotes Treg survival by interacting with Sema4a on Tconv cells(*29*)), while C9-Tregs uniquely express interferon-response genes(*30*) (*Bst2*, *Tgtp2, Ifit1*, and *Isg15*) (Fig. S8, A and B).

Six Treg clusters (C3-C5, C7, C8, and C10) showed an activated phenotype with the lowest expression of lymphoid-tissue homing markers and highest expression of activation genes (Fig. 2D). While activated cell states express high *Tnfrsf9* (encoding for costimulatory activator 4-1BB), each population also expresses distinct genes (Fig. 2E). Differential gene expression of these activated clusters compared with C1-resting cluster or C2-primed/activated cluster or each other was shown (Fig. S6, B to E, Fig. S9, and Fig. S10). Both C4 and C8 Tregs express high *Il1rl1* (ST2) compared to all other states(*2, 31-35*), while they differentially express T-cell activation regulators and effector molecules (C4: *Ass1, Klrg1, Gzmb, and Tagln2 vs.* C8: *Itgb7, Cd52, and GITR*) (Fig. 2E, Fig. S6, B to E, Fig. S9B and Fig. S10B). Similarly, C4 and C8 Tregs expresses high CXCR3, a chemokine receptor regulated by T-bet and associated with tissue Tregs that suppress Th1 responses(*36*) (Fig. 2E). On the other hand, C3 Tregs express genes required for follicular regulatory phenotype (C3: *Maf(37)*) and genes that promote Treg survival and persistence (C3: *Nr4a2, Cst7*) (Fig. 2E and 3C, Fig. S6B and S9A), while C5 Tregs express effectors that promotes suppression (C5: *Il10(38), Gzma(39) and Gzmb(40, 41)*) or tissue protection (C5: *Areg(42)*) (Fig. 2E and 3C, Fig. S6 B to E, and Fig. S9C). C7 Tregs is a small population that expresses stress response genes (*Hspa1a* and *Hspa1b*). C10 Tregs is a small population as well, expressing genes associated with NK cells (*NKG7, CCl5, CXCR6, Gzmb*) and cytokine (*Ifng*), which is expressed in ex-Tregs(*43*) (Fig. S10).

**Fig. 3.**
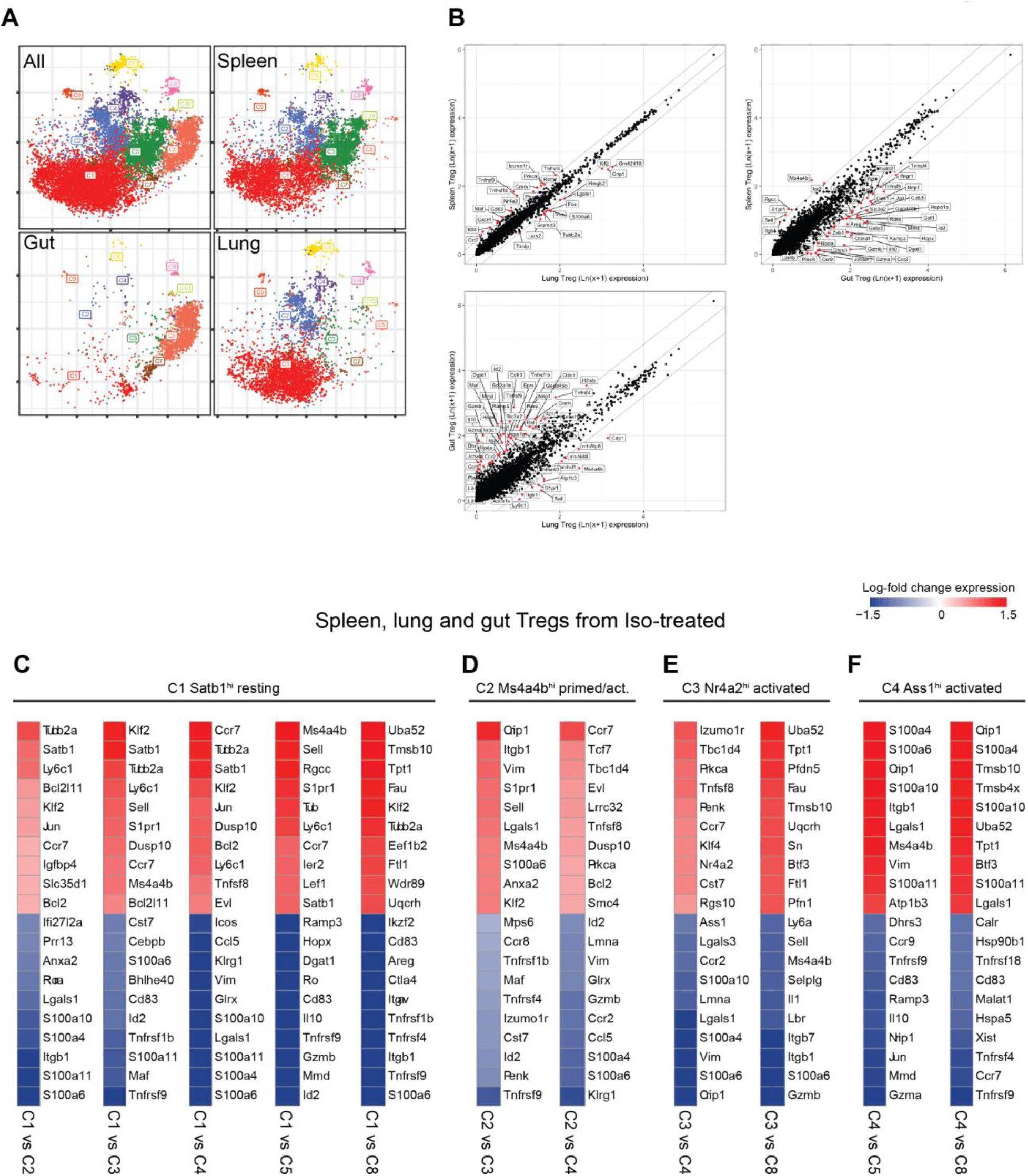
Heterogeneous landscape of Treg cell states defined between the spleen, lung, and gut. **(A)** t-SNE projection of Treg cell states in the spleen, lung, gut, and combined at steady state (Isotype control treated). Individual cells are colored by Treg cell state classification from Figure 2. **(B)** Comparison of Treg signature genes between all tissue-specific cells in the spleen vs lung, spleen vs gut, and lung vs gut at steady state. Significant genes are shown in red (adjusted *p*-value < 0.05, MAST). **(C-F)** Select heatmaps showing the top ten most differentially expressed genes from all Tregs (spleen, lung and gut) of Isotype-control (Iso) treated mice when comparing between two Treg cell states using MAST. Genes are colored by the difference in log-fold-change expression between cell states.

In addition, we identified one proliferative cell state (C6) (*Mki67* and *Top2a*) that co-expresses activation markers (Fig. S9D).

### Analysis of Treg clusters at steady state identified tissue adaptations of Tregs

We next sought to characterize the Treg landscape in the spleen, lung and gut from mice treated with isotype control using the classification of Treg states described in Figure 2 as a reference criterion. At the tissue level, the spleen and lung share a >40% frequency of C1-resting and minor frequencies of primed/activated and activated Tregs. Conversely, >80% of gut Tregs are activated and the majority are C5-Tregs (Fig. 3, A and B). While lymphoid organs, such as the spleen, are known to contain large reservoirs of resting Tregs that can be recruited during immune challenge, the prominence of resting Tregs in the lung is noteworthy and suggests that the lung plays an active role in steady-state immune surveillance.

Activated states of Tregs differ substantially between the tissues, indicative of tissue adaptations acquired by Tregs (Fig. 3A). The spleen in unperturbed state uniquely contained over 30% C3-Nr4a2^hi^ activated Tregs, which express differentiation-promoting transcription factor *Nr4a2*(*44*). In contrast, the lung did not contain the C3-Nr4a2^hi^ Tregs as a main activation state. Rather, the lung contains the C4-Ass1^hi^ and C8-Itgb7^hi^ activated clusters and C6-Stmn1^hi^ proliferative cluster. Comparison of C3 and C4 revealed that C3 cluster differentially express *TBC1D4*, a transcription factor highly expressed in follicular regulatory Tregs, suggesting C3-Nr4a2^+^ activated Tregs include Tfr in the spleen (Fig. 3E). The coexpression of immunomodulatory genes (*Cst7*, *Izumo1r*(*45*), *Nt5e(CD73)*(*46*)) and a reduced activation phenotype (*Tnfrsf9*^med^*Cd83*^med^) without expression of effector proteins compared to other activated states suggest that Nr4a2^+^ Tregs are activated Tregs that are not terminally differentiated but still have suppressive capabilities. Compared with C1-resting Tregs, C4-cluster Tregs express high levels of effector molecules (*GzmB, Glrx, and Klrg1*) and chemokine receptor (*Ccr2*), suggesting this cluster may contain terminally differentiated Tregs with tissue homing receptors(*19*) (Fig. 3, C to F).

The gut Tregs contain small proportion of C1-resting Tregs and very little C2 or C9 prime/activated Tregs. Instead, they primarily contain C5 and C7-activated Treg populations. The C5-IL10^hi^ Tregs uniquely express effectors such as IL10 and tissue repairing Areg (*47*) (Fig. 2E, Fig. S9C) and pro-inflammatory transcriptional regulator *Rora(2)* and elevated levels of gut-homing chemokine receptor *Ccr9(48).* The C7-Dnajb1^hi^ activated Tregs express early response inflammatory (*Dnajb1(49)*, *Hspa1a*(*50, 51*), *Tagap*(*52*), interferon-response-related) genes, some of which are associated with enhanced Treg function and autoimmunity (Fig. S10A). Given that cells in the gut regularly encounter diverse commensal microbes and food antigens, these Tregs may actively promote tolerance even during steady state conditions.

### Reorganization of Treg landscape in multiple tissues after treatment with murine IL-2 mutein

We demonstrated the expansion of Tregs after murine IL-2M *in vivo* (Fig.1, Fig. S1, and Fig. S2), but whether this treatment will expand all Treg sub-populations equally is not clear. Therefore, we compared the frequency of Treg states in each tissue before and after IL-2M to detect changes in molecular phenotypes and function. Treatment with IL-2M shifted the frequency of Treg clusters, reducing C1-resting and C3-activated Tregs while elevating proliferation (C6), primed/activated (C2), and activated Treg states (C4 and C8) (Fig. 4A). These shifts were consistent in all animals (Fig. 4B) and reflected differences at the tissue level, as lung and spleen Tregs shared most of the genes following IL-2M treatment (Fig. S11A), but they were distinct when compared with gut Tregs (Fig. S11, B and C). In the spleen, lung, and gut, C2-primed Tregs and C4-activated Tregs all displayed an over four-fold increase in all animals (Fig. 4C). This expansion of C2- and C4-Tregs in the spleen and lung established these two states as the most prominent Treg populations after IL-2M stimulation, coinciding with a drastic reduction in C1-resting Tregs in both tissues and in C3-activated Tregs in the spleen. A two-fold increase was also observed in C8-activated Tregs in the spleen and lungs. The selective expansion of C4- and C8-Tregs, which both co-express *Tnfrsf9* and *Il1rl1*, suggests that IL-2M may skew expansion in eligible tissues toward specific activation phenotypes. Despite an increase in C2, C4 and C6 Treg states after IL-2M treatment, gut Tregs still maintained C5-activated Tregs as the most prominent cell state (Fig. 4, B and C).

**Fig. 4.**
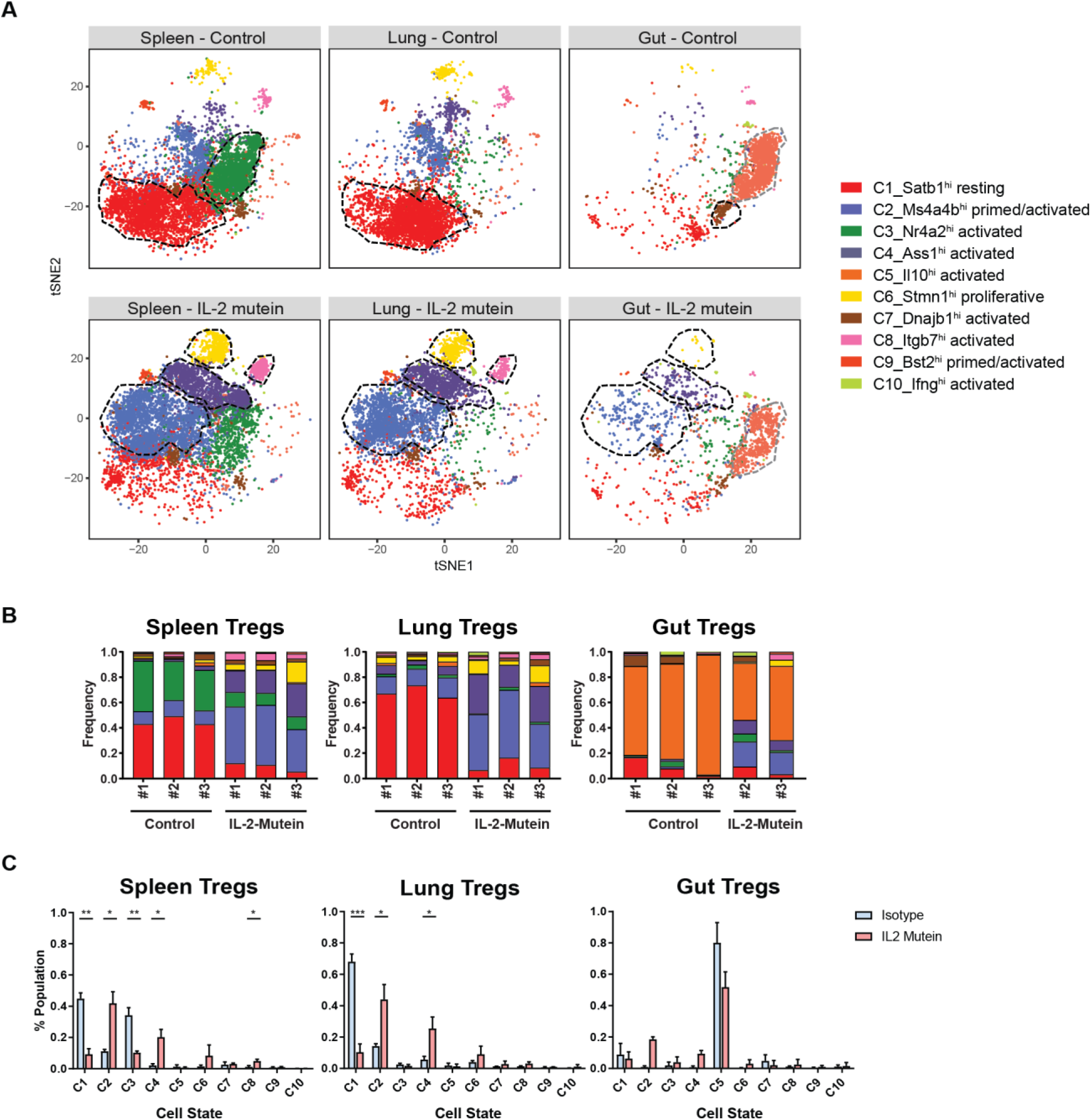
IL-2 mutein induces a convergent expansion toward distinct Treg states in the spleen, lung and gut. **(A)** t-SNE projection of Treg cell states in the spleen, lung, and gut from Isotype control treated (Iso) and IL-2 mutein treated mice. Treg clusters that are enriched at least two-fold in Isotype control group or in IL-2 mutein treated mice relative to the other treatment condition are outlined in with a black dotted line. Prominent populations (>10% of total Tregs in a given tissue) whose frequencies are consistent across treatment are highlighted in gray. **(B)** The tissue-specific frequency of Treg cell states for each experimental replicate. **(C)** Bar plots showing shift in Treg population in the spleen, lung and gut. N=3 for all isotype and IL-2 mutein treatments except for the IL-2 mutein treated gut Tregs (N=2). All comparisons performed using Student’s paired t-test. * *p*<0.05, ** *p*<0.01, *** *p*<0.001

### IL-2 mutein skews clonal Treg expansion toward specific activation states

During development in the thymus, each T cell generates a unique TCRα and -β rearrangement (or clonotype), which predicates antigen specificity and influences Treg fate. This clonotype is retained in cells that descend from a common progenitor T cell, enabling characterization of Tregs that share a developmental lineage. Since we observed an expansion in Tregs (1) toward distinct cell states (C2, C4, and C6) in all three tissues, (2) toward the C8 state in the spleen and lungs, and (3) away from the C3 state in the spleen, we analyzed TCR information combined with Treg cell state classifications from single cells to elucidate whether these cell states expand independently of one another or whether they are developmentally related.

From the 3,600 sequenced single-cell transcriptomes where TCRα and TCRβ were also recovered (see Methods), 3,405 unique Treg clonotypes (1,428 in Iso- and 1,977 in IL-2M-treated conditions) were identified. Of these clonotypes, 119 clonal families (*i.e.* clonotypes that were shared by at least two Tregs) consisting of 314 total Tregs were identified. Among the 119 clonal families, 67 clonal families of Tregs existed in different phenotypically distinct activated states (*e.g.* C3 and C4) or in primed/activated and activated states (*e.g.* C2 and C4), at both steady state and after IL-2M stimulation (Fig. S12), revealing that different Treg cell states can be derived from the same parental Treg. The identification of transcriptional diversity among Tregs from the same clonal family was an interesting result, since we also find that pairs of T cells belonging to the same clonotype tend to be transcriptionally correlated than randomly sampled pairs of Tregs at the population level, although this was a modest effect(*20*) (Fig. S13). Clonal family membership also revealed evidence of Treg migratory patterns between tissues, since we observed that Tregs of the same clonotype spanned the spleen, lung and gut (Fig. S12). Taken together, we interpret these findings as such that, although the TCR responses may generally influence the transcription of Tregs toward a programmed transcriptional fate, clonally-related Tregs still can differentiate into multiple cell states depending on the tissue.

We next characterized cell state classifications of clonotype pairs to identify cell states that may be developmentally linked (which we refer to as “cell state axes”). At steady state, no cell state axis comprised more than 1% of all clonotypes, since the majority of clonotypes (95.6%) did not belong to a clonal family (Fig. 5B, top panel). In contrast, IL-2M-induced clonal proliferation facilitated the detection of prominent cell state axes connecting clonotype pairs by increasing the proportion of Tregs detected in a clonal family (CF) by 105% (211 CF Tregs in IL-2M compared to 103 CF Tregs in Isotype), despite only a 41% increase in Tregs with TCRs recovered (2107 TCRs in IL-2M, 1493 TCRs in Isotype). Multiple axes higher than 1% frequency were identified following IL-2M treatment (Fig. 5B, bottom panel). Among those, multiple clonotypic pairs were observed in the C4-Tregs, including proliferation-dependent C4-C4 (3.9%) and C4-C6 (2.7%) clonotype pairs, as well as C2-C4 (1.0%) and C3-C4 (1.1%) pairs, revealing that C2, C3, and C4 are developmentally linked (Fig. 5, A and B). C4 also shared clonotypes with activated states C7 and C8, albeit at a lower frequency of 0.5% and 0.2%, respectively.

**Fig. 5.**
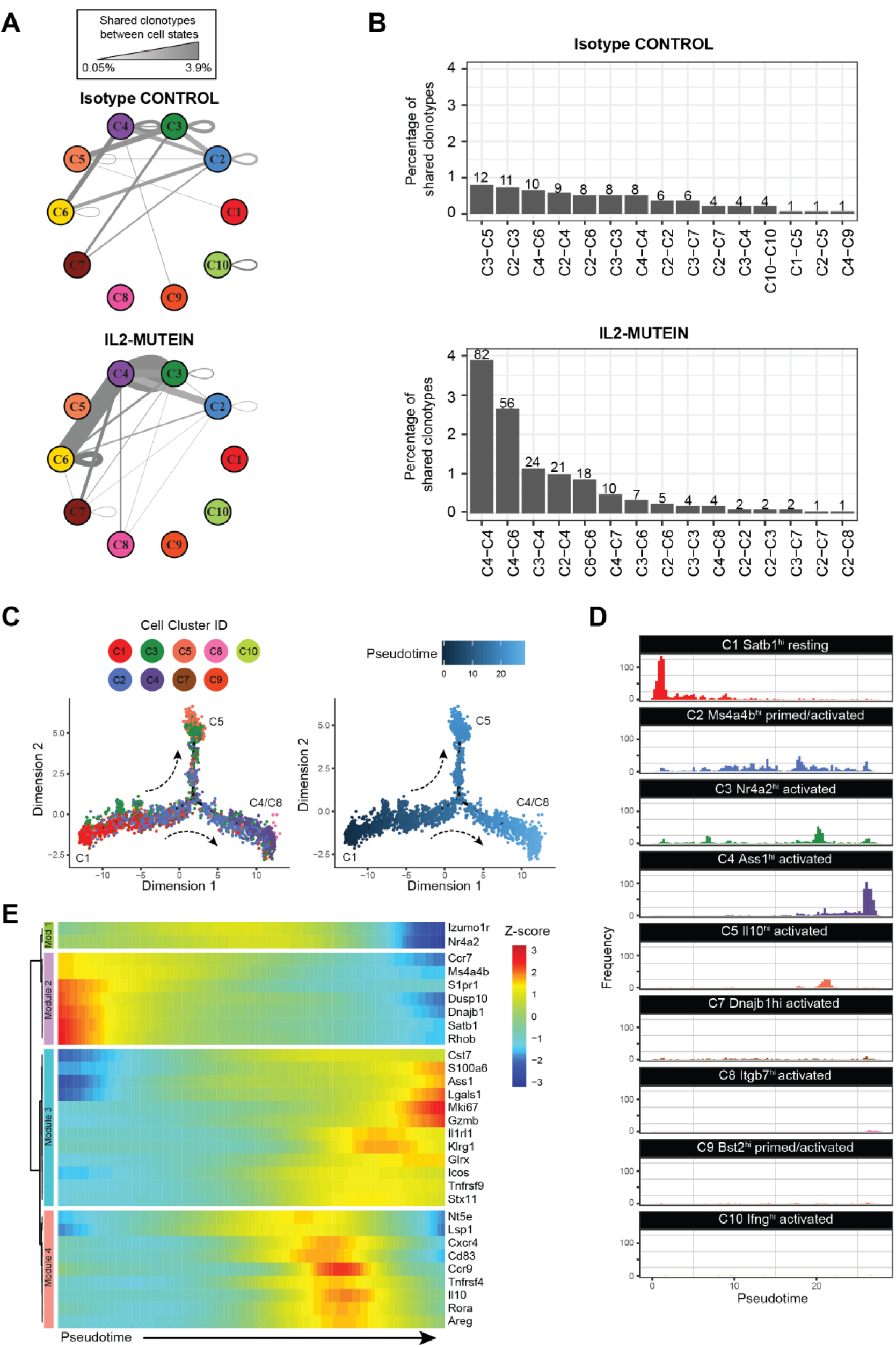
Treg lineage tracing and trajectory analyses identify axes of Treg differentiation across cell states. **(A)** Network graph representing the frequency of shared clonotype pairs (*i.e.* pairs of cells belonging to the same clonotype that exist across two cell states) of spleen and lung Tregs under isotype control- or IL-2M-treated conditions. The thickness and darkness of the gray edges connecting two cell states corresponds to the frequency of shared clonotypes between two cell states or within a single state. **(B)** Top 15 pairs of cell states that share clonal family membership. Numbers above each bar indicate the total number of clonotype pairs in a given cell state axis. **(C)** Pseudotemporal ordering of Tregs with recovered TCRs (*n*=3,600 cells). Individual cells are colored by cell state (left) or by pseudotime, or progress along Treg differentiation (right). **(D)** Histograms showing the distribution of each Treg cell state along the pseudotime axis. **(E)** Heatmap showing the Z-score scaled expression of 30 highly variant genes along pseudotime. Genes are grouped into modules with similar expression by unsupervised hierarchical clustering.

Given that IL-2M increases clonal Treg expansion, we also examined how IL-2M influences the localization of CF Tregs by comparing the frequency of CF Tregs that were shared across tissues versus within the same tissue. While the percentage of inter-tissue CFs remained the same in both Isotype and IL-2M conditions (3.9% versus 4.2%, respectively), the percentage of CFs found only in one tissue was nearly doubled (3.0% versus 5.8%, respectively). Although we cannot ascertain whether *in situ* expansion or migration from other tissues is responsible for this phenomenon, these results indicate that CF Tregs after IL-2M-induced clonal expansion are more frequently observed in the same tissue than across multiple tissues.

### C4 and C8 Tregs expand after IL-2M stimulation through C2/C3 intermediate states

Given the gene expression profiles and robust lineage relationship of the C2-primed states and C3/C4/C8 activated states, we used pseudotime analyses to define their developmental relationship(*53*) (Fig. 5, C and D and Fig. S14A). Treg cell states occupied distinct territories in pseudotime. As expected, C1-resting Tregs occupied earliest period of pseudotime. C2-primed/activated and C3-activated Tregs express markers of previous TCR signaling and were dispersed throughout intermediate points in pseudotime. At the latest periods in pseudotime/differentiation, we observed two distinct differentiation branchpoints consisting of C5-Tregs at one terminus and C4/C8 Tregs at the other (Fig. S14B). This suggests that Tregs generally follow a differentiation trajectory from C1 into either C5 Il10-producing Tregs or C4/C8 *Tnfrsf9^+^Il1rl1^+^* Treg that express effector molecules such as *Klrg1*, *Gzmb, and Glrx*. We obtained similar ordering of cell states when applying pseudotime analyses to the other Tregs in our dataset where no TCR information was obtained (Fig. S14, C and D).

The analysis of Treg trajectories also revealed gene expression dynamics that advance Tregs toward effector states. Of the thirty most variant genes identified from this analysis, four major gene modules were identified that correspond to cell-state classifications (Fig. 5E). Module 1 consisted of *Izumo1r* and *Nr4a2*, the key markers of C3-Tregs, which are moderately expressed all throughout pseudotime until terminal differentiation. The expression pattern of *Nr4a2* agrees with previous findings that establish its role in tonic TCR signaling to allow Treg persistence and survival(*44*). Module 2 consisted of genes most frequently found in resting (C1) and primed/activated (C2/C9) Tregs, which are highly expressed at early cell states and become quickly downregulated once differentiation initiates. Modules 3 and 4 consisted mainly of genes that were highly expressed in the C4/C8 and C5 terminal states, respectively. Overall, analysis of clonal Treg differentiation trajectories suggests IL-2M promotes differentiation into the terminally differentiated C4 and C8 Tregs state by expanding through C2 and C3 intermediate states in the spleen and lung.

### Among IL-2M expanded sub-populations, Tnfrsf9^+^Il1rl1^+^ Tregs demonstrated superiority in in vitro suppression functional assay

Pseudotime analysis demonstrated that bifurcation of Treg differentiation leading into the C4/C8 activated state, showing enrichment of genes in the Module 3 (Fig. 5E). Among those genes, *Il1rl1* and *Tnfrsf9* are highly expressed in the Module 3. Likewise, scRNA-seq demonstrated increase expression of *Il1rl1* and *Tnfrsf9* in the spleen and lung following IL-2M treatment (Fig. 6A). Flow cytometry analysis confirmed that Foxp3^+^ Tregs expressing both Il1rl1(ST2) and Tnfrsf9 (4-1BB) were increased to approximately 10-fold from the lung and spleen of the mice treated with IL-2M at protein level (Fig. 6B). Pan-Tregs isolated after IL-2M treatment increased suppressive activity (Fig. 1 C and D), but it is not clear which sub-state Tregs contribute to this enhanced suppressive activity. Because Tregs with expression of *Il1rl1* and *Tnfrsf9* after IL-2M treatment represent the end stage of the differentiation, we determined suppressive activity of Tregs with differential expression of ST2 and 4-1BB using *in vitro* suppression assays examining suppression activity of Tregs on proliferation and IFNγ production (Fig.6c-d). To this end, Tregs were expanded *in vivo* by murine IL-2M administration. Four days after treatment, four populations of Foxp3^+^ Treg cells based on the expression of ST2 and 4-1BB were sorted and *in vitro* suppression assay was performed. Proliferation of effector T cells was suppressed by Tregs expressing either ST2 or 4-1BB, but ST2^+^4-1BB^+^ Foxp3^+^ Tregs displayed the most superior suppression (Fig. 6C). Similarly, ST2^+^4-1BB^+^ Foxp3^+^ Tregs were most efficient in suppressing IFNγ production of T effector cells under Th1 skewing condition (Fig. 6D). Combined with increased suppression of Foxp3^+^ Tregs expanded with murine IL-2M in Fig.1, this data demonstrated that murine IL-2 mutein treatment preferentially increased the proportion of ST2^+^4-1BB^+^ Foxp3^+^ Tregs, which possess superior ability to suppress proliferation and IFNγ production of T effector cells. Thus, endogenous expansion of Tregs using *in vivo* murine IL-2M administration has the potential either to induce or expand Foxp3^+^ Tregs with the highest capacity to suppress autoreactive T cells.

**Fig. 6.**
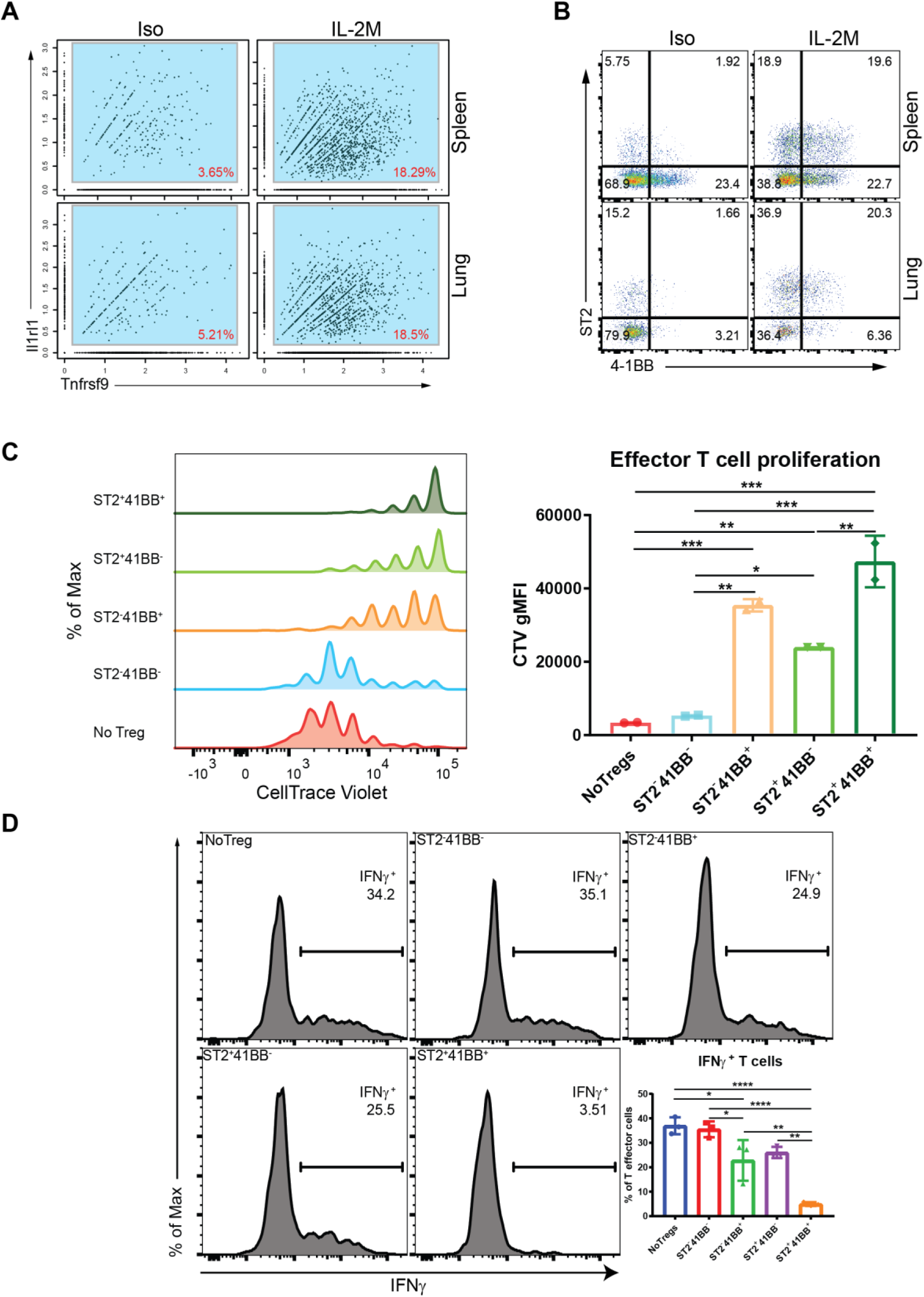
Tnfrsf9^+^Il1rl1^+^ Tregs are superior in suppression functional assay of Tregs *in vitro*. (**A**) Expression of *Il1rl1(ST2)* and *Tnfrsf9(4-1BB)* in RNA single cell analysis of Tregs from the spleen (*top*) or lung (*bottom*) of Isotype control-treated (Iso) or murine IL-2M-treated mice (IL-2M). (**B**) Expression of ST2 and 4-1BB using FACS analysis from the spleen (*top*) or lung (*bottom*) of Iso or IL-2M-treated mice. (**C**) Fopx3-EGFP+ Tregs were stained based on ST2 and 4-1BB expression. Four quadrants of Fopx3-EGFP+ Tregs were sorted based on ST2 and 4-1BB. *In vitro* suppression assays were performed using four populations of sorted Tregs or without Tregs. Dilution of CTV fluorescence intensity (*left*) was shown as CTV-labeled naïve T cells proliferated and diluted fluorescence intensity. (**D**) Suppression function of four Treg sub-populations on IFNγ production. Four quadrants of Fopx3-EGFP+ Tregs were sorted based on ST2 and 4-1BB. Tregs were cocultured with effector T cells and APCs differentiated under Th1 skewing condition. IFNγ intracellular staining under the conditions using Tregs from PBS- or IL-2M treated mice. No Tregs were added in one condition as a control. *Statistics*: **C-D**, One-way ANOVA for multiple comparisons. **0.001< *p* <0.01, ***0.0001< *p* <0.001, \*\*\*\**p* < 0.00001

## Discussion

IL-2 plays an indispensable role in immune tolerance by governing the proliferation and maintenance of Foxp3^+^ Tregs. IL-2 GWAS studies identified multiple polymorphisms affecting IL2 pathway genes encoding IL2Rα (CD25) and IL-2 as key genetic risk alleles in autoimmunity, corroborating IL2 as an essential cytokine for maintaining immune homeostasis(*54–59*). Together with IL-2 therapy studies and mouse models defective in IL2 signaling, it is now well-accepted that IL2 is the key cytokine necessary for generation, function, and survival of Foxp3^+^ Tregs. Consistent with genetic studies in human and mouse, reduction in numbers and functions of Foxp3^+^ Tregs had been reported as a contributing factor for a variety of autoimmune diseases(*59–65*).

Given its role in Foxp3^+^ Treg biology, providing IL2 to Foxp3^+^ Tregs as an intervention for autoimmune diseases had been tested in clinic over the years. Recent clinical trials using low-dose recombinant human IL2 (Proleukin^TM^, aldeskeukin) have shown preliminary success in selectively expanding Tregs in chronic graft-versus-host-disease (cGVHD)(*7, 10, 66*) and in steroid-refractory moderate-to-severe systemic lupus erythematosus (SLE) patients(*67*). Additional efforts have focused on novel forms of IL2 with superior pharmacokinetics and Treg selectivity. These include a long-lived bivalent IgG-IL2 fusion protein with N88D mutation that reduces IL2Rβ binding(*12*), a PEG-modified murine IL2 that increases IL2 retention in the body by protection from enzymatic digestion and renal clearance(*68*), and a murine IL2/anti-IL2 antibody (JES6-1) conjugate(*69*). Despite the effort to develop various forms of IL2 with superior pharmacokinetics and selectivity for treatment of autoimmune and inflammatory diseases, it is not clear how they affect heterogeneity of Tregs, given that Tregs exists in a variety of sub-states at different tissue sites, that are phenotypically and functionally heterogenous.

To establish an effective Treg-mediated therapy, it is critical to characterize the phenotypic and functional heterogeneity induced by the treatment and define mechanisms that promote functional Treg differentiation. Due to the diverse profiles of proteins expressed in Tregs, it is challenging to define functional Tregs, let alone to identify Tregs with superior effector functions. CD4^+^CD45RA^low^CD25^hi^CD127^low^ cells have been previously used as human surface Treg markers to identify the pan-Tregs, and various activated markers of Tregs have been adopted to understand the functional and phenotypic diversity of Tregs. However, the detailed molecular characterization of Treg cell states is limited with a mere handful of activation markers. Single-cell RNAseq overcomes this obstacle by providing a comprehensive view of transcriptional heterogeneity of Tregs, thereby making it possible better design a targeted Treg therapy.

Using scRNA-seq and *ex vivo* cellular assays, we mapped the landscape of Treg sub-states at the single cell level during steady state and following murine IL-2M treatment in mice. We used an engineered, half-life extended murine IL2 mutant protein (IL-2M) and demonstrated that it expanded Tregs robustly *in vivo.* Expanded Tregs by murine IL-2M treatment suppressed proliferation of Tconv as well as production of IFNγ from Th1 effector cells. Single cell RNA-seq study uncovered that IL-2M induces a convergent shift of the Treg landscape toward primed (C2) and highly suppressive activated Treg states (C4, C8) in addition to proliferative Treg state (C6). Furthermore, TCR-seq combined with scRNA-seq revealed a migratory network across tissues. IL-2M-mediated Treg expansion increased shared TCR clonality and this expansion was more prominent in clonal families that were identified in one tissue, while the percentage of inter-tissue clonal families remained the same in both Isotype and IL-2M treatment. Interestingly, shared TCR clones were also found in multiple and distinct cell states, providing us with a molecular snapshot of the transcriptional changes following IL-2M stimulation.

Cell clustering of Treg single-cell data generated a universal criterion of Treg states to detect variation in the molecular profiles of Tregs across tissues and treatment. Many of the marker genes, which have previously been studied in Tregs as a bulk population, allowed us to infer functions associated with Treg sub-populations. As a result, cell clustering identified ten clusters with distinct combinatorial transcriptional profiles, which could be more generally grouped into one resting, two primed-activated, one proliferative, and six activated cell subsets. The signature genes that classified these populations align well with previous Treg-focused scRNA-seq studies(*4, 20*). For example*, S1pr1, Ccr7, Sell*, and *Satb1* identify resting Tregs, while suppressive Tregs express genes such as *Il10, Rora*, and *Tnfrsf9*. Our study also identified primed-activated populations, and showed that they share a developmental lineage, express intermediate levels of resting and activated genes, and can give rise to suppressive (C5) or terminally differentiated (C4/C8) effector Tregs. As our TCR analyses suggest, we believe these clusters should be considered as activation states along the Treg differentiation continuum and not Treg subsets that arise independently.

The similarities in single cell profiles between the spleen and lung versus the gut at steady state suggest that the spleen and lung in healthy mice possess a large proportion of resting-Tregs (C1-resting Tregs) that serve as a reservoir of Tregs in preparation for potential inflammatory stimuli. The spleen uniquely contains a high frequency of activated Tregs (C3-activated Tregs) expressing Nr4a2, a Foxp3-binding transcription factor (*70*), Cst7(*71*), a marker for previous Treg activation and regulator of cytotoxicity, and Izumo1r(*45*), a marker for natural Tregs in the spleen. Compared with other activated Treg cluster C4, C3-activated cluster also differentially express *Tbc1d4* and *Maf*, transcription factors for follicular regulatory T cells(*37, 72*). Given the role of the spleen in immune surveillance and its lymphatic connection to peripheral tissues as well as germinal center formation(*73*), these splenic Nr4a2^+^ Tregs may include follicular regulatory T cells (Tfr) and recently activated Tregs licensed to exit the spleen and migrate into sites of inflammation. On the other hand, the gut is composed of uniquely activated Treg clusters, IL10-producing suppressive C5 and inflammatory/stress response-associated C7 clusters, likely due to its constant exposure to commensal microbes, pathogens and food antigens, it requires many activated Tregs that can maintain tissue homeostasis and promote tolerance(*74, 75*).

IL-2M treatment reduced the proportion of resting C1 Tregs in the spleen and lungs and the C3 Tregs in the spleen, while expanding the numbers of proliferating (C6), primed/activated Tregs (C2) and potently suppressive Tnfrsf9^+^Il1rl1^+^ activated Tregs (C4 and C8). The same C2, C4, and C8 populations were also expanded in the gut, although the Il10-producing C5 Tregs remained as the most prominent population the gut. Since we observed an over two-fold expansion in the number of Tregs in all tissues following IL-2M stimulation, the decrease in C1 and C3 Treg percentage was likely driven more by the concomitant expansion of various expanding sub-populations as opposed to a significant reduction in the absolute numbers of the prominent Treg populations during steady state. Both expanded activated Treg populations, C4 and C8, express high Il1rl1 (ST2), which is the receptor for tissue alarmin IL-33(*33*). ST2^+^ Tregs have been previously shown to prevent excessive tissue damage in organs such as the skin, liver, and gut(*2, 76, 77*). C4 and C8 Tregs, while both expressing Tnfrsf9 and Il1rl1, as well as effector molecules such as *Klrg1* and *Gzmb*, differ in expression of regulatory mediators such as *Ass1*, *Cd52* and *Tnfrsf18*. Trajectory analysis suggests that the C4 and C8 Tregs are terminally differentiated. Previously, it has been shown that Klrg1^+^ Tregs represent a terminally differentiated Treg subset that is recently activated by antigen and resides in the lamina propria of small intestine(*19*). Klrg1^+^ Tregs are highly activated and express enhanced levels of suppressive Treg molecules. Interestingly, the development of Klrg1^+^ Tregs requires extensive IL-2R signaling. Based on gene expression, it is highly likely that C4 and C8 Tregs contain Klrg1^+^ Tregs. Between the two clusters, the C8 appears to be more terminally differentiated due to its expression of effector cell surface receptors Cd52 and Tnfrsf18-encoded GITR(*35, 78*), which suppress other immune cells by binding to ITIM-containing Siglec10 and GITR-L, respectively. Identification of these expanded clusters suggests that IL-2M may significantly remodel the Treg landscape towards more terminally differentiated effector Tregs and suppressive Tregs, which mediate potent immunosuppression in these tissues.

We provide evidence using TCR analyses and trajectory analyses to suggest that C4 and C8 Tregs differentiate from C2/C3 primed/activated states. TCR analyses identify that C2, C3, C4, and C8 Tregs in the spleen and lung share clonotypes, demonstrating that they can all derive from a common progenitor. Cells from these populations that share the same clonotype are also present across the spleen and the lung, revealing an immune trafficking axis that exists between the two tissues. Using trajectory analysis to order cells along a differentiation continuum, we show that C2 and C3 Tregs contain cells throughout the trajectory manifold, suggesting these sub-states are transitional state between resting and further differentiated or activated Treg states. C4/C8 Tregs, on the other hand, are mostly present on one of the termini on the trajectory manifold. Taken together with the shared clonotype information, this suggests that IL-2M-expanded C4 and C8 populations come from C2 and C3 populations. The ordering of identified cluster-specific marker genes correlates with classical genes known to identify resting (i.e. *S1pr1 and Ccr7*) and activated Tregs (i.e. *Gzmb, Klrg1*, and *Icos*), suggesting that the trajectory manifold recapitulates patterns of Treg differentiation. Trajectory analysis also identifies a bifurcation in Treg differentiation after IL-2M treatment, which either differentiate into suppressive Il10^+^Rora^+^ C5 Tregs, which are most prevalent in the gut, or into C4/C8 Tregs that are prominent in the spleen and lungs.

In this study, we characterize Tregs in the spleen, lung, and gut using scRNA-seq and analyze how the Treg landscape in these tissues is influenced by IL-2M stimulation. We identified an immune axis of clonal TCRs between the spleen and the lung and found an activated, Tnfrsf9^+^Il1rl1^+^ Treg populations that expand highly upon IL-2M stimulation. Supporting analyses indicate that this population differentiates through multiple primed and recently activated intermediate cell states and it is a highly suppressive Treg state that inhibits Th1 effector cell activity *ex vivo*. Overall, this study reveals a potential mechanism by which IL-2M induces tissue-specific Treg immune modulation and classifies Treg state signatures that may serve as biomarkers for studying Treg responses. While our analysis focused on genes with previously characterized immune cell functions, previously uncharacterized genes that were associated with Treg states of differentiation, such as certain ribosomal genes, warrant further investigation to elucidate their role in Treg biology.

## Materials and Methods

### Mice and IL-2 mutein treatment

All experimental studies were conducted under protocols approved by the Institutional Animal Care and Use Committee of Amgen. Animals were housed at Association for Assessment and Accreditation of Laboratory Animal Care International-accredited facilities at Amgen in ventilated micro-isolator housing on corncob bedding. Animals had access ad libitum to sterile pelleted food and reverse osmosis-purified water and were maintained on a 12:12 hour light:dark cycle with access to environmental enrichment opportunities. Foxp3EGFP (#006772), C57BL/6J and B6.SJL-PrprcaPepcb/BoyJ (BoyJ) mice (8-12 week old) were purchased from Jackson Laboratory. Once purchased, mice were housed under specific pathogen-free conditions in the laboratory animal facility and were handled according to protocols approved by IACUC at Amgen. In order to determine the effect of murine IL-2 mutein (IL-2M), different amounts of IL-2M (0.1, 0.33, and 1 mg/kg) were administered into naïve C57BL/6J mice. After four days of IL-2M treatment, flow cytometry was performed to determine its effect on the regulatory and conventional T cells. In order to determine the effect of murine IL-2M on T cell proliferation, Ki67 staining was performed. Untreated, PBS- or mouse IgG Fc isotype (ThermoFisher Scientific, 31205) – treated mice were compared and used as negative controls. Administration of murine IL-2M via intraperitoneal (i.p.) or subcutaneous (s.c.) routes resulted in similar effects. For scRNA-seq experiment, mice were administered with murine IL-2 mutein (0.33 mg/kg) or mouse IgG Fc isotype control (Iso) by intraperitoneal (IP) injection and were sacrificed 4 days after murine IL-2M administration.

### IL-2/anti-IL2 antibody complex (IL-2C)

IL-2 complex (IL-2C) was prepared as described(*79*) by incubating 1 μg recombinant mouse IL-2 (PeproTech) with 5 μg purified anti-mouse IL-2 antibody (JES6-1, BioXcell) for 30 min at 37°C (molar ratio for IL2:anti-IL-2 is 2:1). IL-2C (total 6 μg) was administered i.p. in a final volume of 200 μl daily for three days. Four days after the last administration of IL-2C, expansion of Tregs and Tconv was determined by FACS analysis.

### Flow cytometry reagents and Treg sorting

Anti-CD11c (N418), anti-F4/80 (BM8), anti-CD19 (6D5), anti-CD11b (M1/70) and anti-CD45.1 (A20) were from Biolegend. Anti-ST2 (U29-93), anti-CD3 (17A2), anti-CD4 (RM4-5), anti-IFNγ (XMG1.2) and anti-CD16/32 (2.4G2) were from BD Biosciences. CellTrace Violet (CTV), Fixable Viability Dye eFluor™ 780 and Foxp3 (FJK-16s) were from Fisher Scientific. Anti-4-1BB (158332) was from R&D.

### Isolation of murine spleen and lung Treg cells for single-cell RNA seq

The spleen and lung were collected and processed according to the protocol from Miltenyi murine Lung Dissociation kit (130-095-927). Lungs were perfused via the right ventricle before the process. Briefly, tissues were cut and rinsed in PBS. Transfer tissues into the gentleMACS C Tube containing enzyme mix of Enzyme D and A. Run gentleMACS Program m_lung_01 twice and incubate samples for 30 mins at 37 C under continuous shaking at 150 rpm. Run the gentleMACS Program m_lung_02. Filter the tissue disaggregate through 70 um strainers and wash the cells with PEB buffer twice. The samples were ready for downstream flow staining and sorting.

### Isolation of murine gut Treg cells for single-cell RNA seq

The small intestine (extending from the end of stomach to the cecum) was collected and processed according to the protocol from Miltenyi murine Lamina Propria Dissociation kit (130-097-410). Briefly, the feces and Peyer’s patches were removed, and the small intestine was filleted and cut into pieces. The samples were washed in predigestion solution, vortexed and passed through 100 um fliters twice. Then the samples were washed in HBSS (w/o), vortexed and passed through 100 um fliters. The samples were transferred to gentleMACS C Tube containing pre-heated 2.35 mL of digestion solution with Enzyme D, R and A. Incubate the samples for 30 mins at 37 C under continuous shaking at 150 rpm. Plug the C Tube into the gentalMACS and run Program m_intestine_01 twice. Filter the tissue disaggregates through 100 um strainers and wash the cells with PB buffer twice. The samples were ready for downstream flow staining and sorting.

### Treg suppression assay

Various populations of Treg cells were FACS sorted on Aria II sorter (BD Biosciences) based on their expression of EGFP reporter. Naïve CD4+ T cells were isolated by Miltenyi mouse naive CD4+ T cell isolation kit and were used as effector T cells. APCs were prepared by depleting CD90+ T cells from splenocytes and irradiated for 3000 rads (FaxiTron). Naïve CD4+ T cells and APCs were from BoyJ mice which is CD45.1+ to differentiate from EGFP+ Treg cells which were in B6 background by flow cytometry. To test the suppression function of Treg cells on proliferation, various populations of Treg cells (1×10^4^/well) were cocultured with effector T cells (2×10^4^/well) and APCs (3×10^4^/well), as indicated, along with anti-CD3 (BioXcel, 145-2C11, 1 μg/ml) for 72 h. To test the suppression function of Treg cells on IFNγ production, various populations of Treg cells (2500/well) were cocultured with effector T cells (2×10^4^/well) and APCs (3×10^4^/well), along with anti-CD3 (1ug/mL), anti-IL-4 (BioXcell, 11B11, 10ug/mL), IL-12 (Peprotech, 6ng/mL) for 96 h. Then the whole culture will be stained for intracellular cytokine staining (ICS). Briefly, cells were washed once and were stimulated with cell stimulation cocktail (eBioscience) for 5 hours in cell culture medium, then fixed and stained for IFNγ.

#### Single-Cell RNA-seq

Single-cell RNA-seq was performed in three replicates, with each replicate containing the spleen, lung, and gut Tregs from one isotype treated mouse (Iso) and one IL2-mutein treated mouse (IL-2M). Combining different treatment conditions and tissues in each replicate allowed us to identify and regress out gene expression differences driven by batch processing.

CD4^+^Foxp3^+^ single cells were sorted from tissue-dissociated single cell suspensions after gating on CD14^−^CD19^−^CD11c^−^F4/80^−^ cells). After sorting, cells were immediately washed and resuspended in chilled PBS + 0.04% BSA at 100-1000 cells per microliter. Washed cells were then encapsulated in one lane of a 10x Chromium Single Cell Controller, and libraries were constructed using either the 10x Single Cell 3’ Reagent Kit (V2 chemistry) (https://support.10xgenomics.com/single-cell-gene-expression) or the 10x Single Cell 5’ Reagent Kit (https://support.10xgenomics.com/single-cell-vdj) according to the manufacturer’s protocols. Completed libraries were sequenced on the HiSeq 4000 or NovaSeq 6000 platforms (26/8/0/98 Read1/i7/i5/Read2). Data were analyzed using the Cell Ranger 2.0.0 Pipeline by aligning reads to a genome containing mm10 genome and eGFP.

##### Post-alignment filtering and data aggregation

The Seurat implementation(*21*) in Scanpy(*80*) was used to aggregate data from multiple experimental replicates and cluster Tregs into distinct states. Cells with low quality transcriptomes (<500 detected genes) and doublets (>8000 genes) were removed from the analysis. Canonical correlation analysis was performed to identify common sources of variation and align data from different library preparation batches (each batch consisted of one isotype and one IL-2M treated sample of Tregs from each tissue), thereby mitigating batch effects. After filtering, our dataset detected 18,367 total genes and 31,908 cells across all replicates (12,840 cells in replicate 1, 11,822 cells in replicate 2, and 7,246 cells in replicate 3). We recovered an average of 5,032 unique mRNA molecules after collapsing duplicate UMIs and 1,635 genes per cell.

##### Unsupervised cell clustering and differential expression analysis of clusters

Unsupervised cell clustering of scRNA-seq data was performed using the FindClusters() function (resolution = 0.6) from Seurat. Differential expression testing of each cluster (*i.e.* Treg cell state) versus all other clusters was performed using MAST(*81*) to generate a list of differentially expressed genes for each cluster. A list of genes that were significantly upregulated genes by both tests were then used to define hallmark marker genes for each cell state. Hallmark genes for each of the Treg sub-populations was determined by selecting from among the top ten genes with an adjusted p-value < 0.01, average log2-fold change expression > 0.3, and statistically significant enrichment in less than half of the total cell clusters. The inclusion of genes significantly enriched in multiple clusters was set under the assumption that while the combinatorial marker gene expression is unique to each cluster, individual marker genes may be upregulated by multiple cell states. Mitochondrial and pseudogenes were excluded from consideration as gene markers. T-stochastic neighbor embedding using RunTSNE() was utilized from Seurat for dimensionality-reduced visualization of single cell data.

##### Single Cell TCR-seq

For samples prepared using the 5’ Reagent Kit, 2 ul of amplified cDNA was set aside to amplify mouse TCRa and TCRb chains in two sequential target enrichment PCRs using custom reverse primers: 5’-TGAAGATATCTTGGCAGGTG-3’ and 5’-TGCTCAGGCAGTAGCTATAATTGCT-3’ were used for the 1^st^ PCR, and 5’-GATCTTTTAACTGGTACACA-3’ and 5’-TTTGATGGCTCAAACAAGGA-3’ were used for the 2^nd^ PCR. All forward primers, primer concentrations, and PCR conditions and cycles were the same as those specified in the manufacturer’s protocol. Completed libraries were sequenced on the HiSeq4000 platform (150/8/0/150 Read1/i7/i5/Read2). Data were analyzed using the Cell Ranger 2.0 Pipeline by aligning reads to all IMGT mouse TCR variable and constant region sequences(*82*). Germline gene segments and CDR3 TCRa and TCRb sequences were identified and clonotypes were defined by single cells containing a recovered TCRa and TCRb with identical germline gene segments and CDR3 sequences. In total, 3,600 cells contained complete TCR and transcriptomic information out of 7,246 TCRs recovered using the 10x Chromium 5’-VDJ method.

#### Defining cell state networks using TCR clonotype and transcriptomic data

Custom R scripts were used to define relationships between TCR and gene expression information. Single cell TCR data was merged with transcriptomic data using the common cell nucleotide barcode incorporated during reverse transcription. All clonotype analysis was performed using cells with one single productive TCRa and TCR chain and transcriptomic information (filtered using Scanpy), while all other cells were excluded.

To generate chord diagrams to illustrate the clonal relationships and cell states shared by Tregs from the same clonotype, the circlize package in R was used. Tissue of isolation and cell state definitions determined by Seurat were used to construct the outer and inner tracks of the chord diagram. Links were drawn for any two cells sharing a clonotype.

To generate network plots depicting the magnitude of clonotype sharing across cell states, the igraph package in R was used in Fig. 5a. The frequency of clonotypes across cell states was first calculated by summing the number of cell pairs shared between two cell states and dividing by the total number of clonotypes. This value was then used to determine edge widths connecting two cell states, or vertices. Only spleen and lung Tregs were included in this analysis, as we recovered low numbers of gut Treg TCRs.

#### Pseudotime analysis of Treg differentiation

Monocle2 (2.8.0) was used to order cells according to pseudotime. To generate the input data for the analysis, we combined cells from IL-2M and isotype conditions, since (1) cell state classifications were determined independently of treatment and (2) we reasoned that the developmental trajectory should be preserved across treatment condition. We removed cells (i.e. C6 Tregs) whose primary marker genes were driven by proliferative status and not by their relative status along Treg-specific differentiation, although an independent analysis including these cells revealed that they are dispersed evenly throughout the trajectory manifold, which is expected since both resting and activated cell states are capable of proliferation. The gene list used for pseudotime analysis was filtered to include genes expressed in greater than 10% of all cells in the data (5,311 genes). We then used Monocle DDRTree to perform dimension reduction and manifold constructing, representing the trajectory of Treg differentiation. Cells were then ordered along the manifold.

## Acknowledgements

We thank Olga Pryshchep, Hyun Ra, Nathan Deer, Jennifer Cheung, Laura Dieu, Aaron Fojas, Marisela Killian, Min-Zu Wu, Dev Bhatt, Oliver Homann and Oh-Kyu Yoon for their technical support and intellectual input.

