## Supplemental Figures and Figure legends for "Dynamic changes in the regulatory T cell heterogeneity and function by murine IL-2 mutein"

**List of Supplementary Figures:**

Figure S1-Figure S14

Fig. S1

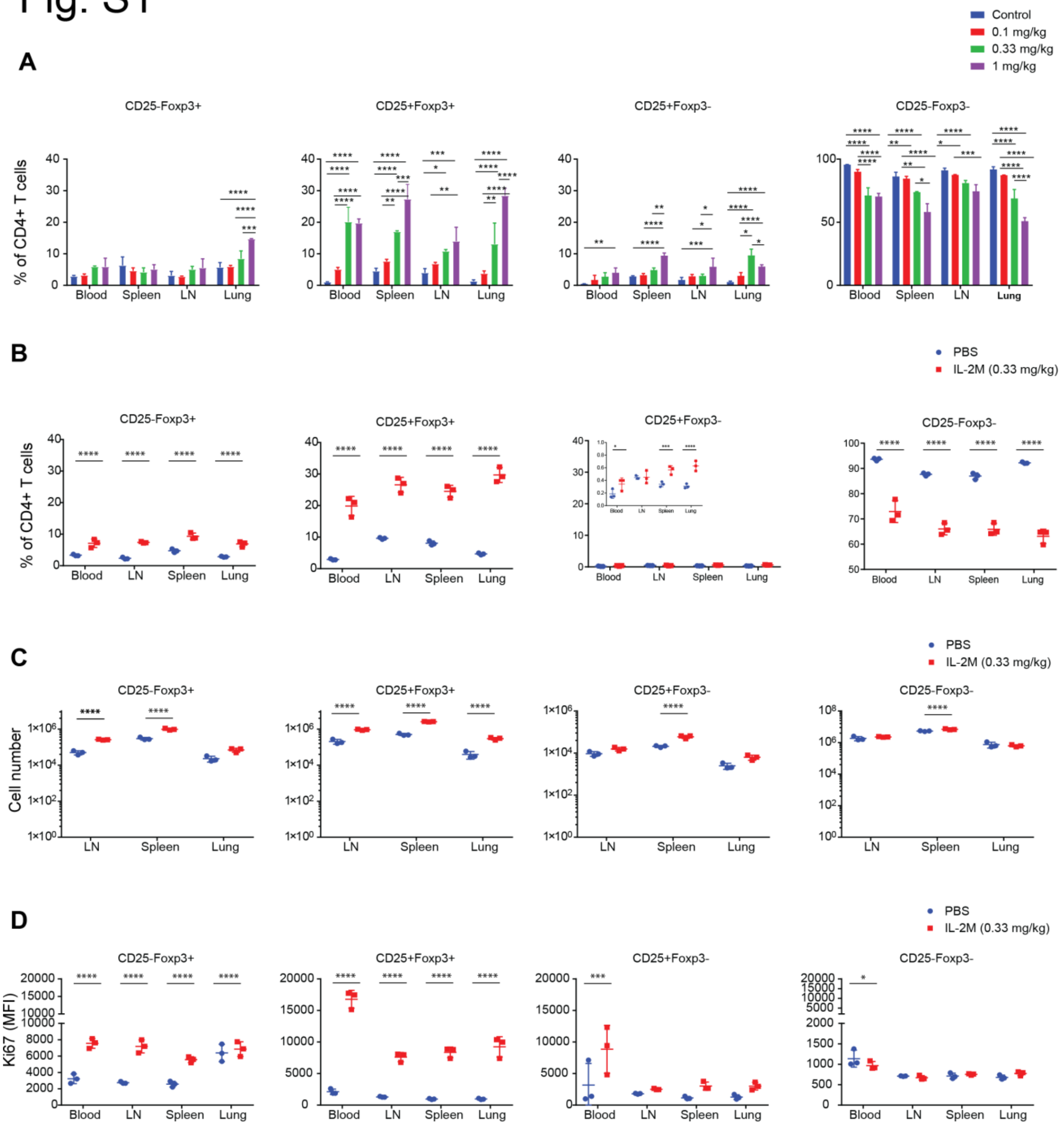


**Fig. S1: Effects of murine IL-2M in expanding mouse CD4+CD25+Foxp3+ Tregs**

**(A)** Dose-dependent effects of murine IL-2M on the percentages of CD25-Foxp3+ and CD25+Foxp3+ Tregs and CD25+Foxp3- activated Tconv and CD25-Foxp3- Tconv within CD4 T cells in the blood, spleen, lymph nodes and lung. Experiments were performed using 3 mice per each group. Control was untreated mice. (**B**) Effects of murine IL-2M (0.33 mg/kg) on the percentages of each population within CD4 T cells from the blood, spleen, lymph nodes and lung. (**C**) Effects of murine IL-2M (0.33 mg/kg) on the cell numbers of each population from the blood, spleen, lymph nodes and lung. (**D**) Effects of murine IL-2M (0.33 mg/kg) on the proliferation of each cell population assessed by expression of Ki67 (mean fluorescence intensity). **B**-**D**, results are representative of at least two independent experiment. Experiments were performed using 3 mice per each group. Control: PBS-treated mice. *Statistics*; Two-way ANOVA for multiple comparisons. *$0.01 < p < 0.05$, **$0.001 < p < 0.01$, ***$0.0001 < p < 0.001$, ****$p < 0.00001$

Fig. S2

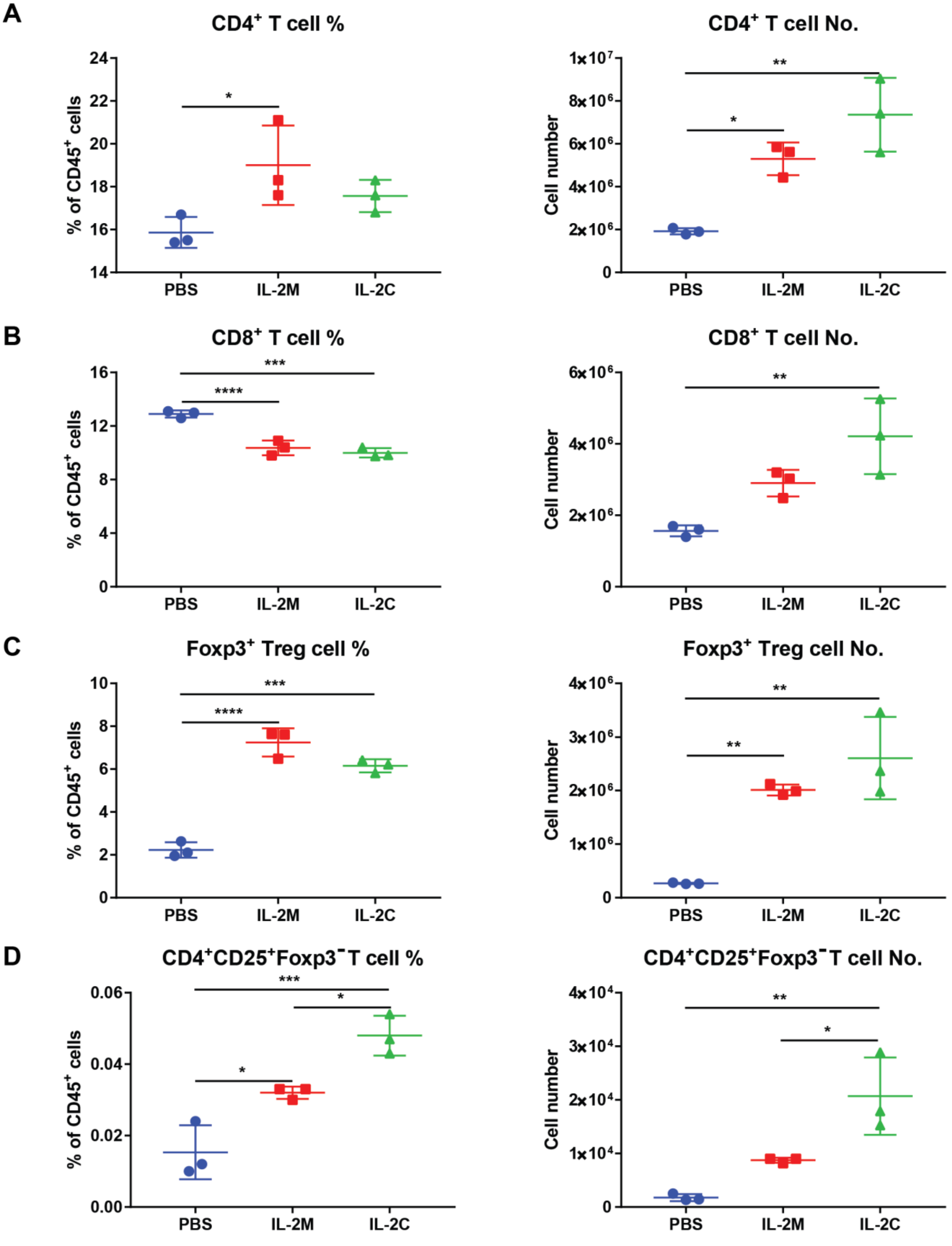


**Fig. S2: Comparing murine IL-2M and IL-2/anti-IL-2 antibody(JES6-1) complex (IL-2C).**

(**A**) Effects of IL-2M and IL-2C on the percentages and cell numbers of CD4 T cells. Percentages of CD4 T cells (PBS vs IL-2M, *: $p = 0.0465$); Cell numbers of CD4 T cells (PBS vs IL-2M, *: $p = 0.021$; PBS vs IL-2C, **: $p = 0.0021$) (**B**) Effects of IL-2M and IL-2C on the percentages and cell numbers of CD8 T cells. Percentages of CD8 T cells (PBS vs IL-2M, ***: $p = 0.0007$; PBS vs IL-2C, ***: $p = 0.0003$); Cell numbers of CD8 T cells (PBS vs IL-2C, **: $p = 0.0061$) (**C**) Effects of IL-2M and IL-2C on the percentages and cell numbers of Foxp3+Tregs. Percentages of Foxp3+Tregs (PBS vs IL-2M, ****: $p < 0.0001$; PBS vs IL-2C, ***: $p = 0.0001$); Cell numbers of Foxp3+Tregs (PBS vs IL-2M, **: $p = 0.0074$; PBS vs IL-2C, **: $p = 0.0017$) (**D**) Effects of IL-2M and IL-2C on the percentages and cell numbers of $CD4^+CD25^+Foxp3^-$ activated Tconv. Percentages of $CD4^+CD25^+Foxp3^-$ activated Tconv (PBS vs IL-2M, *: $p = 0.0235$; PBS vs IL-2C, ***: $p = 0.0009$; IL-2M vs IL-2C, *: $p = 0.0280$); Cell numbers of $CD4^+CD25^+Foxp3^-$ activated Tconv (PBS vs IL-2C, **: $p = 0.0035$; IL-2M vs IL-2C, *: $p = 0.0297$). *Statistics*; One-way ANOVA for multiple comparisons.

### Fig. S3

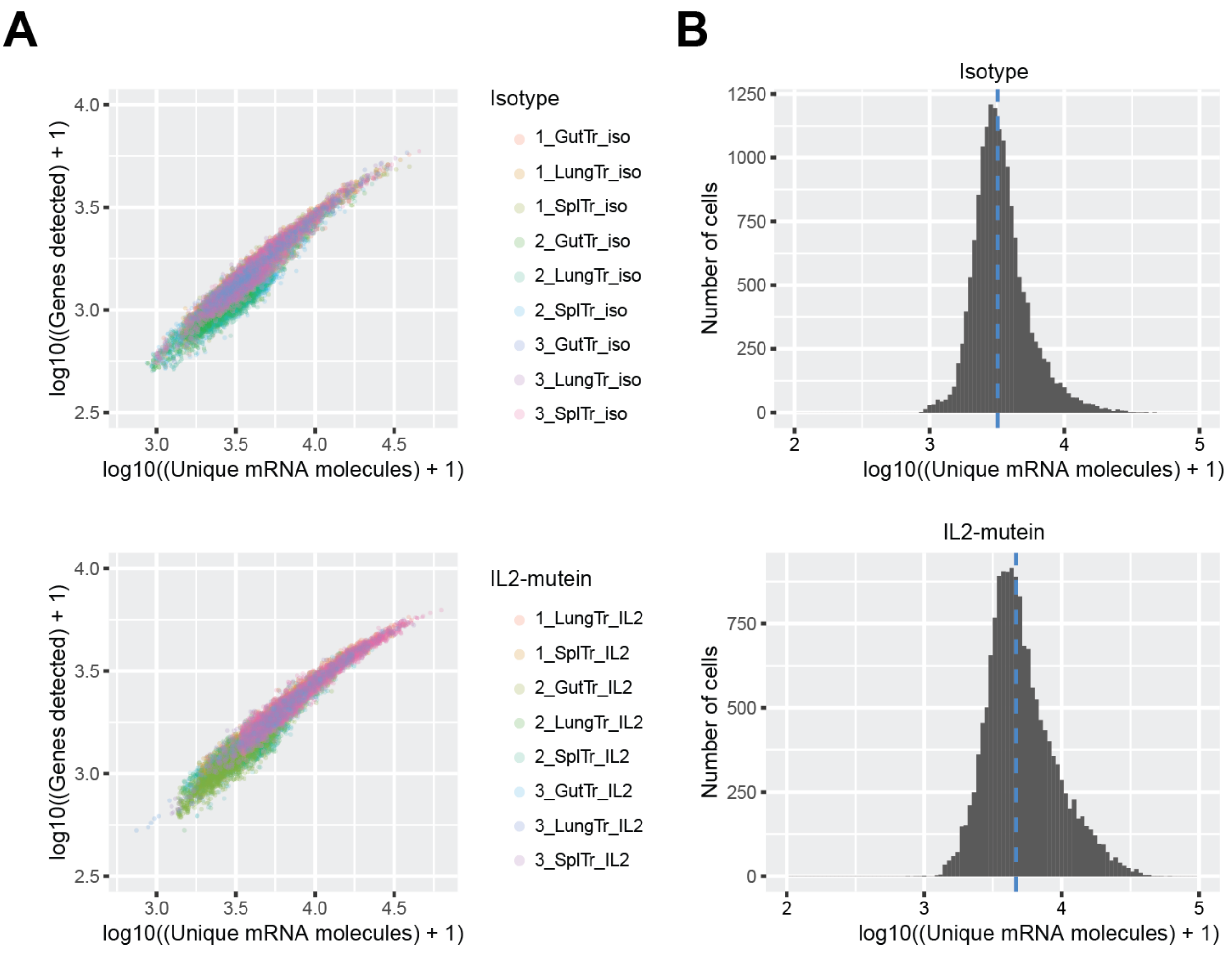


**Fig. S3. Single cell RNA-seq variation across samples and treatment.**

**(A)** Number of genes and unique mRNA molecules for all single cell RNA-seq libraries after isotype control or IL-2M treatment. Single cell libraries are colored by sample. **(B)** Histogram of the distribution of unique mRNA molecules across all single cell RNA-seq libraries. Blue dotted line indicates the mean number of molecules per cell for isotype control and IL-2M treated cells.

Fig. S4

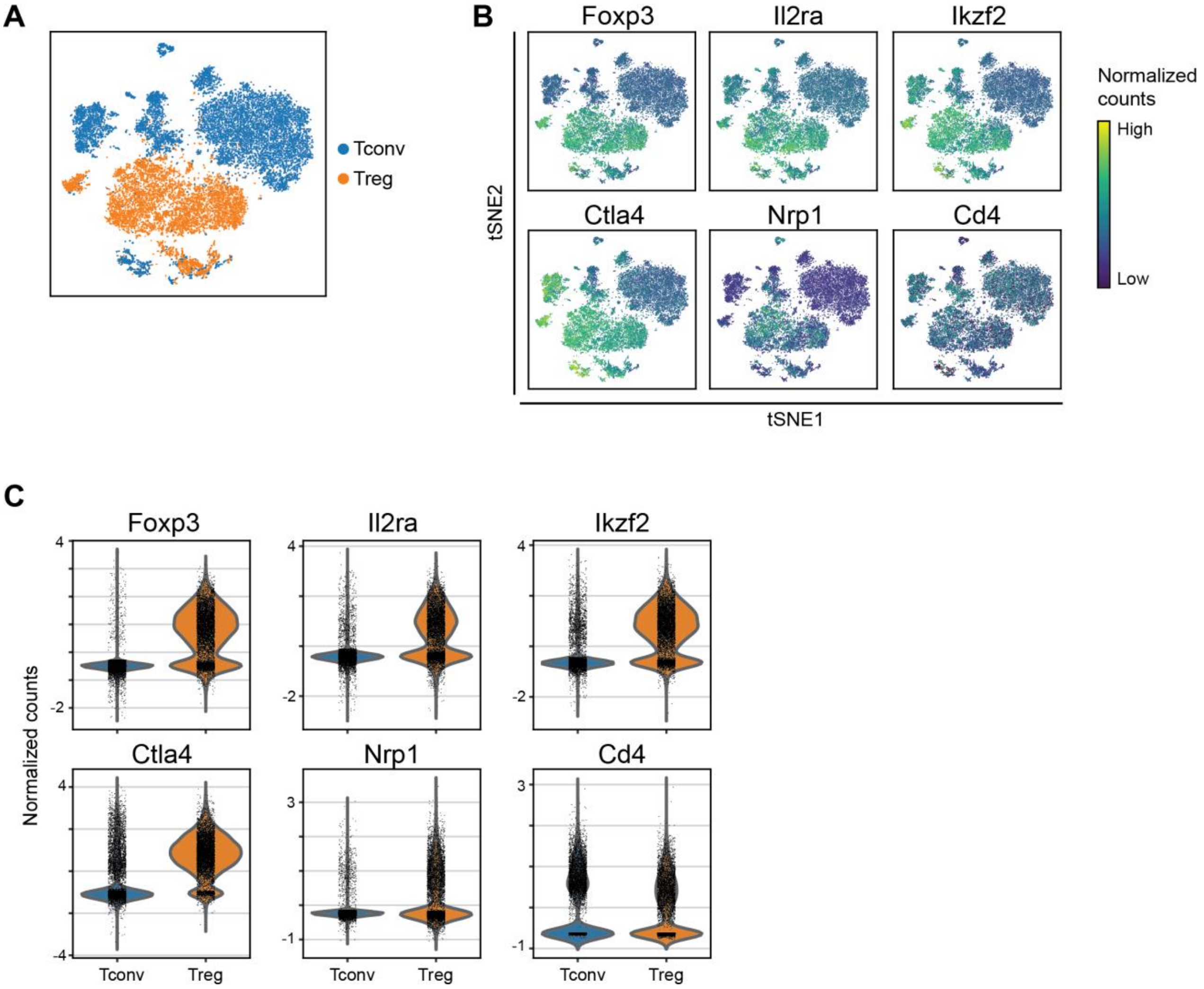


**Fig. S4. Treg expression of established Treg genes relative to Tconv cells.**

**(A)** t-SNE projection of single regulatory T cells (Treg) and conventional CD4 T cells (Tconv). **(B and C)** t-SNE projections **(B)** and violin plots **(C)** show the single-cell expression levels of established Treg marker genes *Foxp3*, *Il2ra*, *Ctla4*, *Ikzf2*, and *Nrp1*, as well as even *Cd4* expression between Treg and Tconv cells.

### Fig. S5

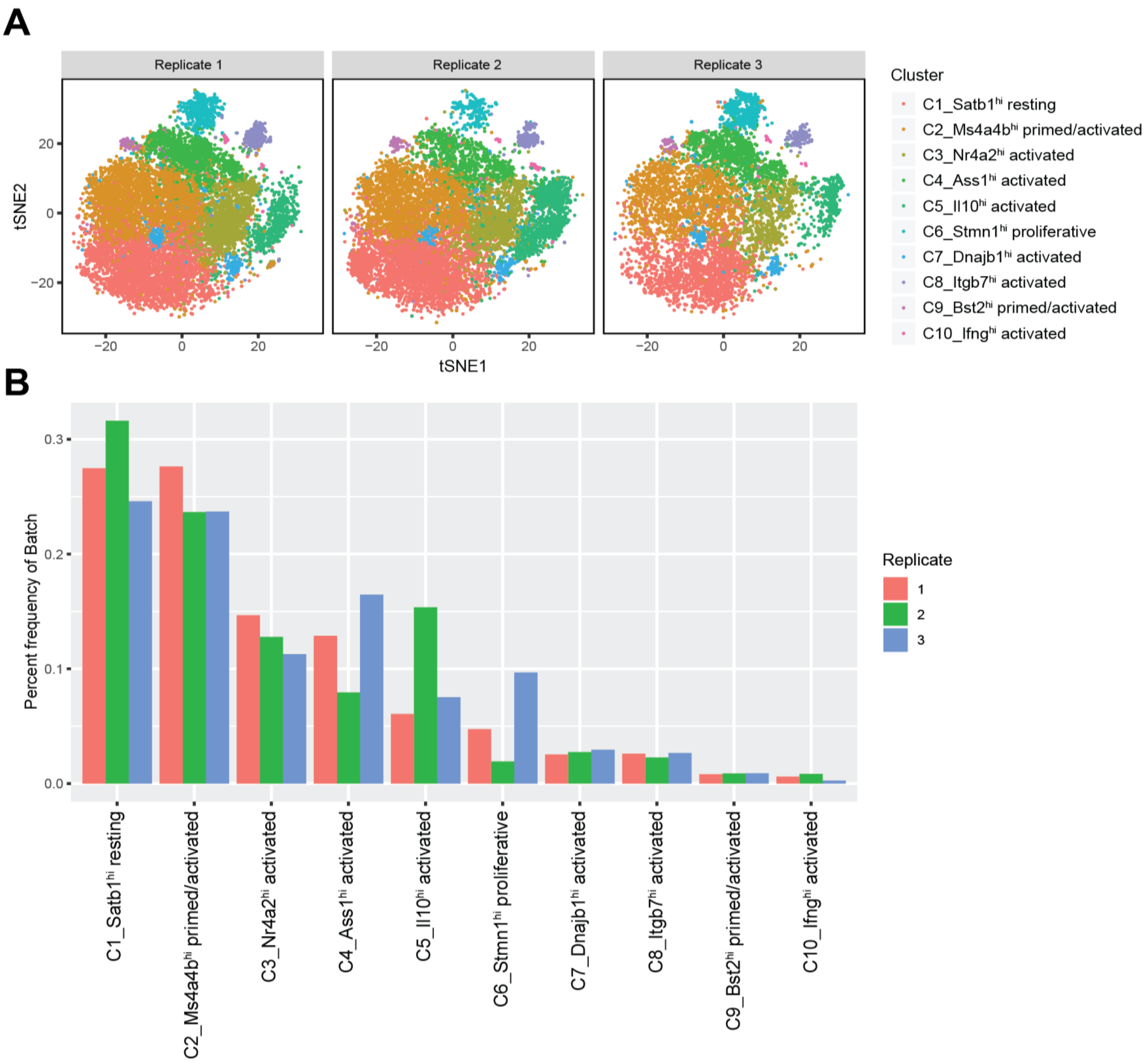


**Fig. S5. Variation of single cell RNA-seq cell state classification across experimental replicates. (A)** t-SNE projection consisting of the cells for each experimental replicate (a.k.a. one isotype treated mouse and one IL2-mutein treated mouse) from which Tregs were isolated and profiled using single-cell RNA-seq. **(B)** Frequency of Treg single cell libraries per experimental replicate assigned to each to Treg cell state classification.

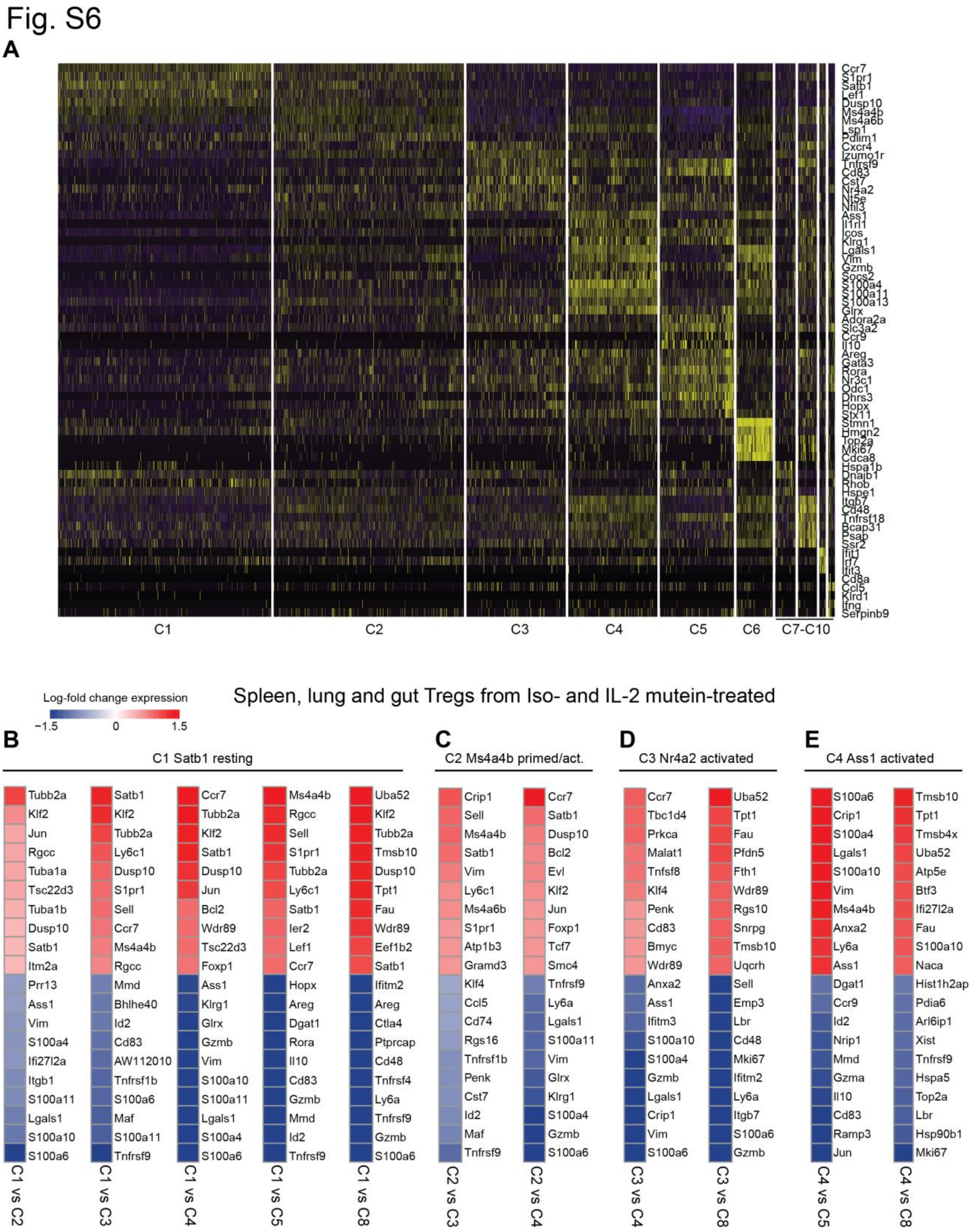


**Fig. S6. Distinct gene expression profiles between Treg states.**

**(A)** Heatmap showing top ten overlapping genes for each Treg cell state following differential expression using Wilcoxon Rank Sum test. Vertical lines depict individual cells belonging to each cell state. **(B-E)** Select heatmaps showing the top ten most differentially expressed genes from all Tregs (spleen, lung and gut) of both isotype control- and IL-2M- treated mice when comparing between two Treg cell states using MAST. Genes are colored by the difference in log-fold-change expression between cell states.

Fig. S7

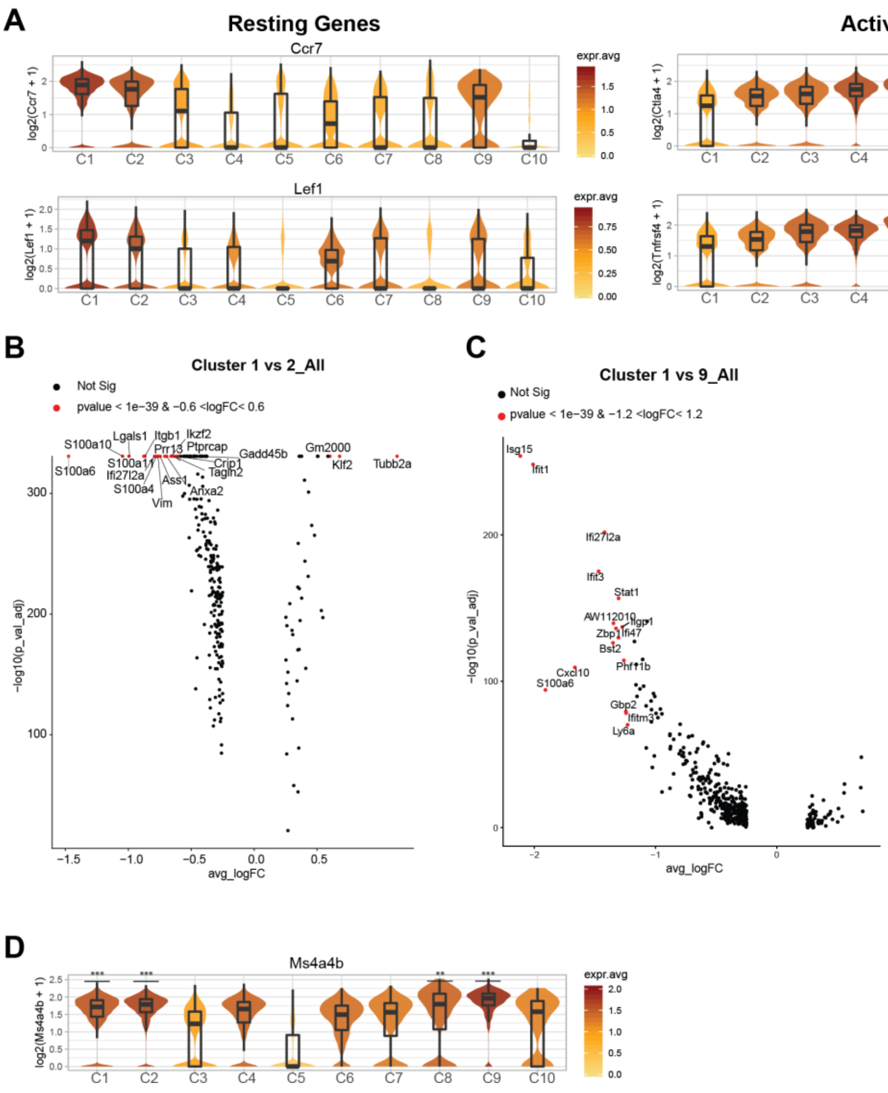


**Fig. S7. Cellular heterogeneity within primed-early activated Treg sub-populations.** **(A)** Violin plots showing intermediate C2 and C9 expression of classical resting and activated Treg marker genes. **(B)** Volcano plot showing gene expression profile between C1 resting and C2 primed/activated Treg states. Labeled genes show most significant genes based on p-value and fold-change thresholds. **(C)** Volcano plot showing gene expression profile between C1 resting and C9 primed/activated Treg states. Labeled genes show most significant genes based on p-value and fold-change thresholds. **(D)** Violin plot showing high Ms4a4b expression primed-early activated Treg states. Clusters with significantly higher gene expression relative to all other cell states are shown with asterisks. ** = P<0.01, *** = P<0.001

Fig. S8

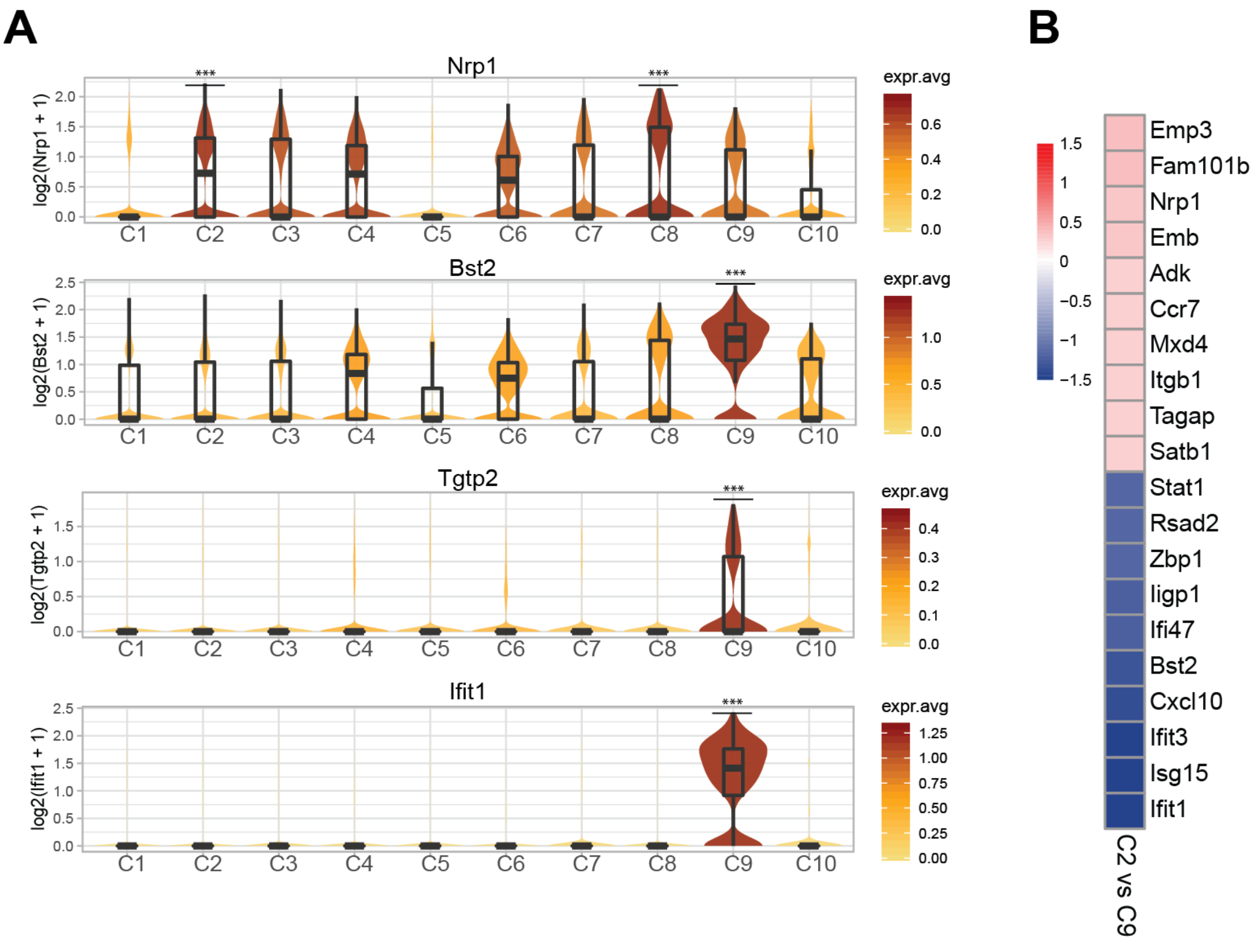


**Fig. S8. Distinct interferon and Treg stability signatures differentiate primed/activated Treg sub-populations.**

**(A)** Violin plots depicting selected signature genes differentially expressed between C2 and C9 primed-activated Treg states. Clusters with significantly higher gene expression relative to all other cell states are shown with asterisks. **(B)** Heatmap showing the top ten most differentially expressed genes between C2 and C9 Tregs using MAST, colored by log-fold-change expression. ** = $P<0.01$, *** = $P<0.001$

Fig. S9

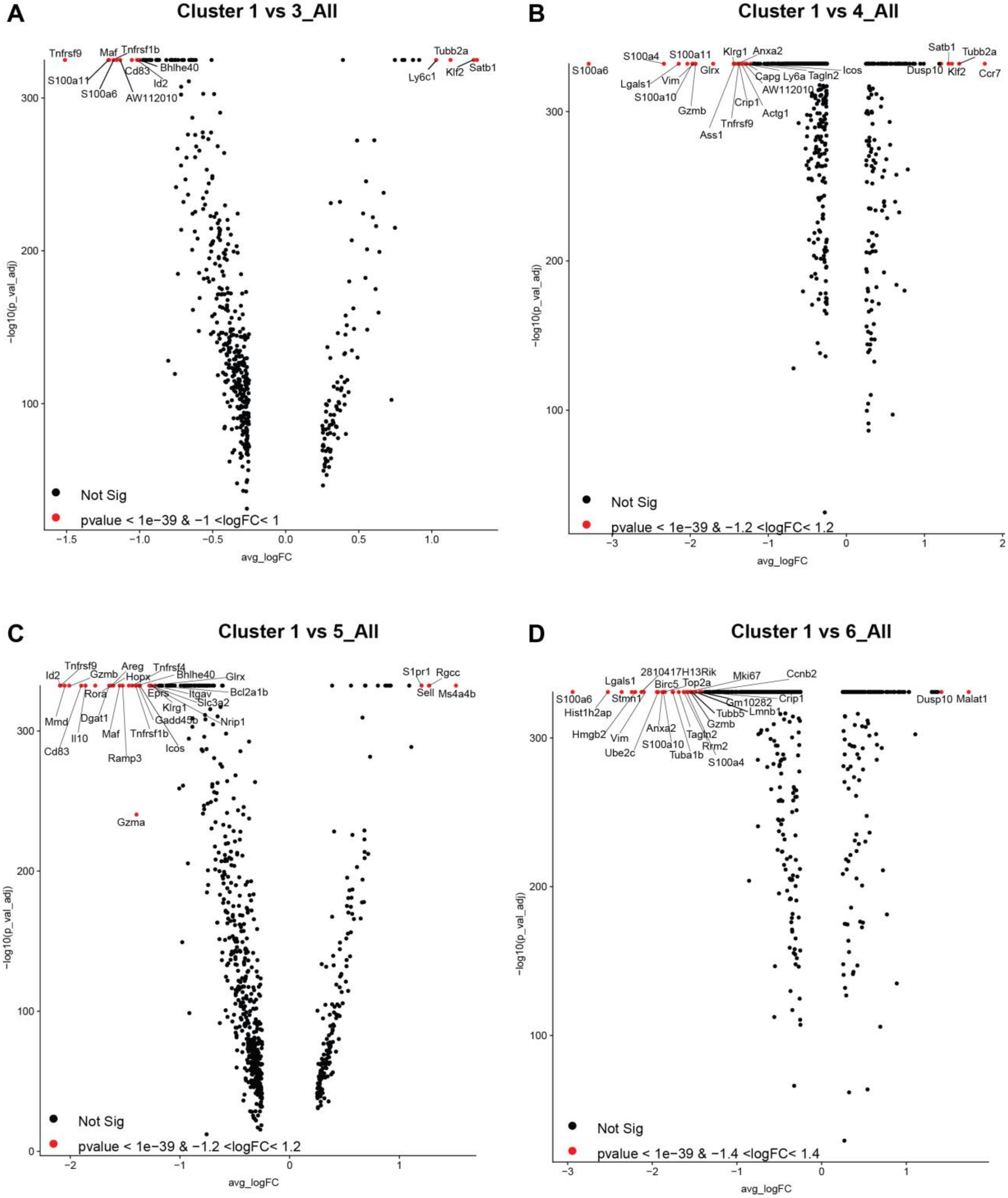


**Fig. S9. Different gene expression profiles between clusters.**

**(A-D)** Volcano plots showing gene expression profiles between C1 Tregs and C3, C4, C5, and C6 Tregs. Labeled genes show most significant genes based on p-value and fold-change thresholds.

Fig. S10

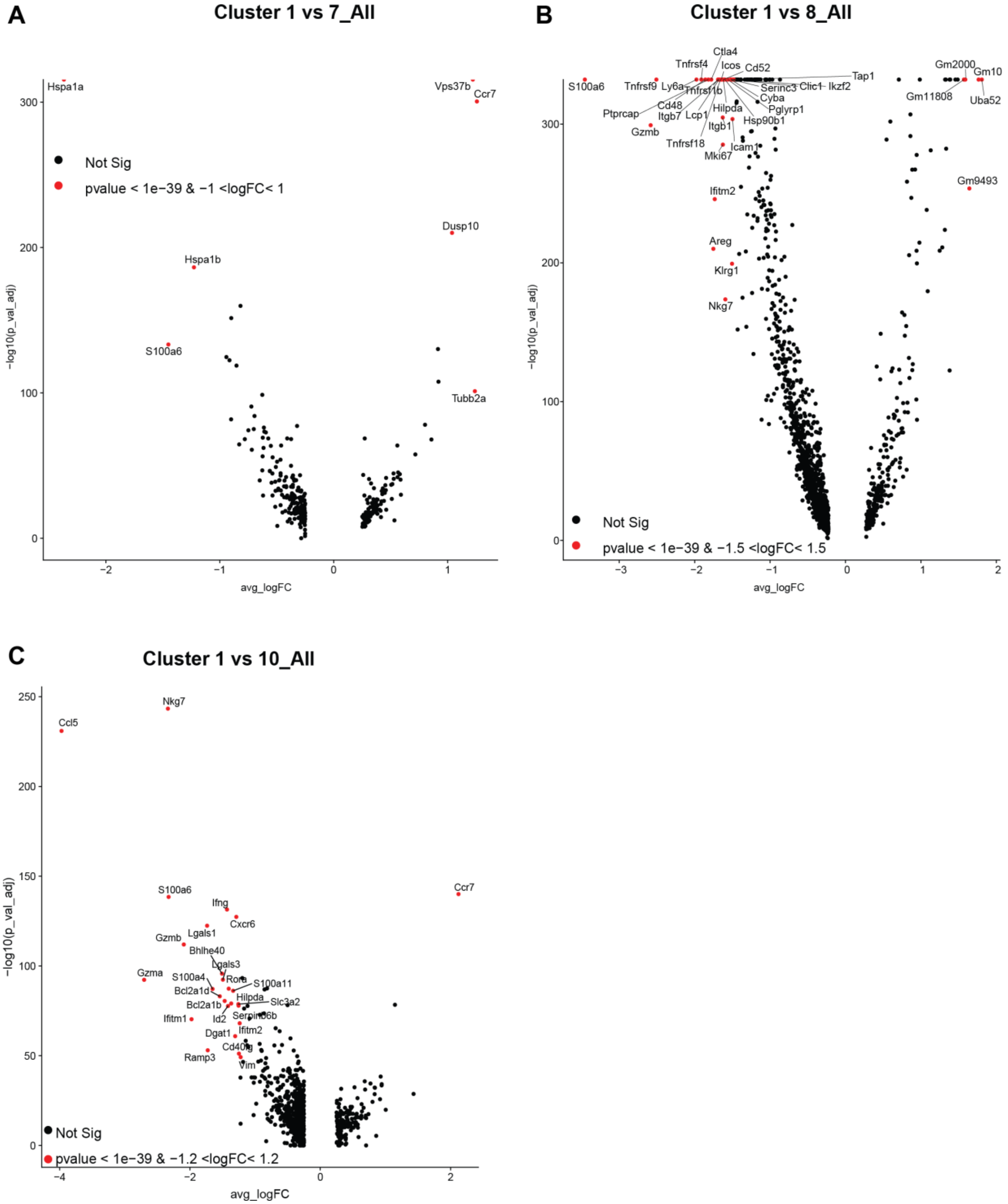


**Fig. S10. Different gene expression profiles between clusters.**

**(A-C)** Volcano plots showing gene expression profiles between C1 Tregs and C7, C8, and C10 Tregs. Labeled genes show most significant genes based on p-value and fold-change thresholds.

Fig. S11

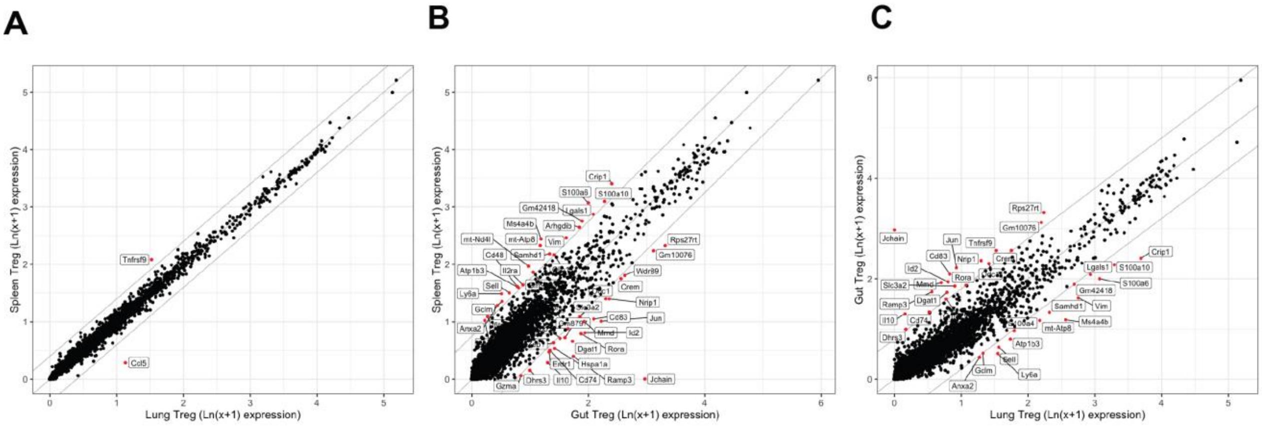


**Fig. S11. Transcriptional profiles converge in the spleen and lung after IL2-mutein treatment, while gut retains a distinct transcriptional profile from the spleen and lung.** **(A)** Comparison of Treg signature genes between all Tregs in the (**A**) spleen vs lung, (**B**) spleen vs gut, and (**C**) lung vs gut after IL-2M treatment. Significant genes are shown in red (adjusted p-value < 0.05, MAST).

Fig. S12

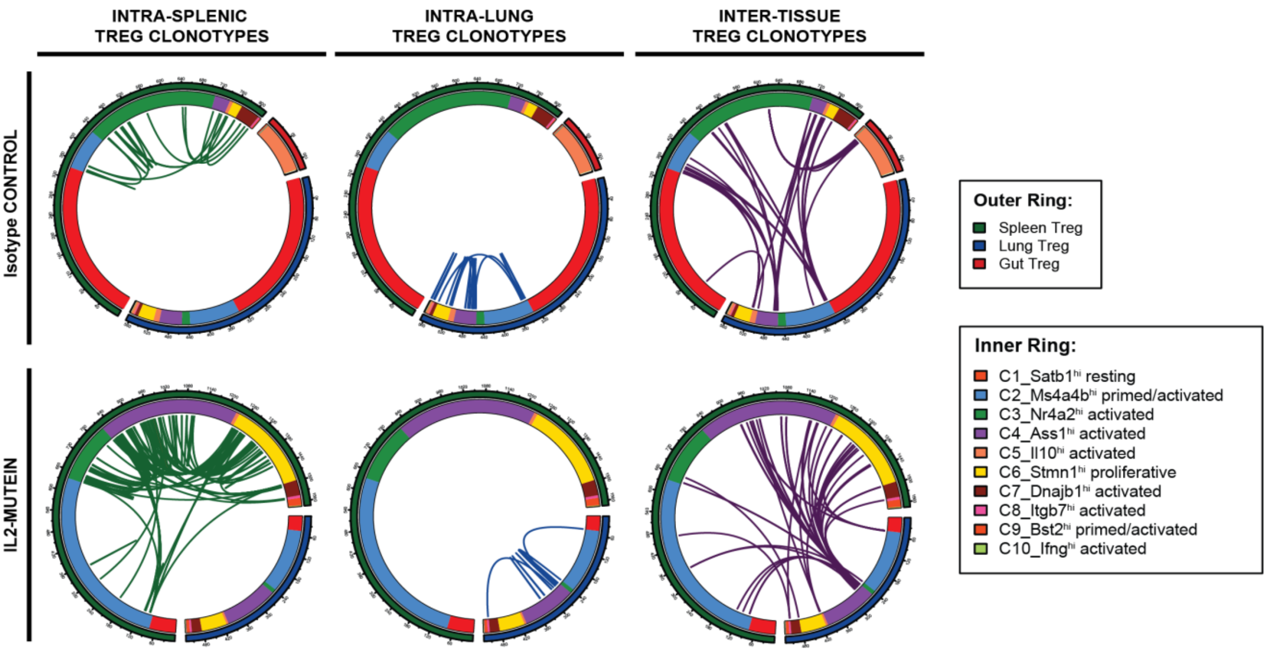


**Fig. S12. Tregs from the same progenitor differentiate across multiple states within and across tissues.**

Chord diagrams of TCR clonotype sharing involving Tregs within the spleen (left), lung (middle), and across tissues (right) in control and IL-2M-treated condition. Outer ring denotes the tissue from which a Treg was isolated. Inner ring denotes the cell state identified by Seurat cell clustering. Green lines link splenic Tregs of the same clonotype. Blue lines link lung Tregs of the same clonotype. Purple lines link clonotypes shared between different tissues.

### Fig. S13

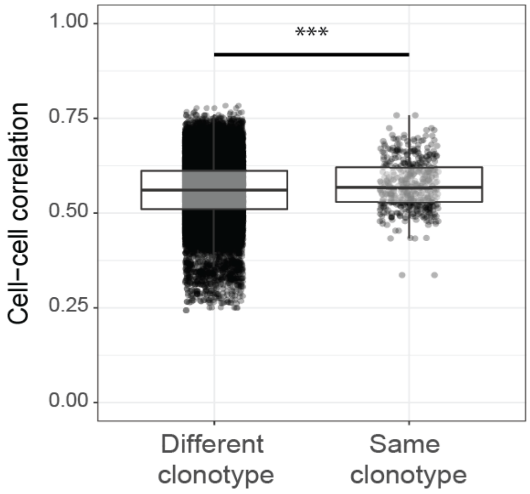


**Fig. S13. Tregs belonging to the same clonotype share modest yet significant transcriptional programs.**
Cell-cell correlation analysis between the transcriptomes of Treg cells that express the same rearranged TCRαβ (n=478 cell-cell correlations) or unrelated TCRs (n=10,000 correlations between any Tregs in the data not sharing clonal family membership, randomly sampled). *** P < 0.0001 (two-tailed t test).

Fig. S14

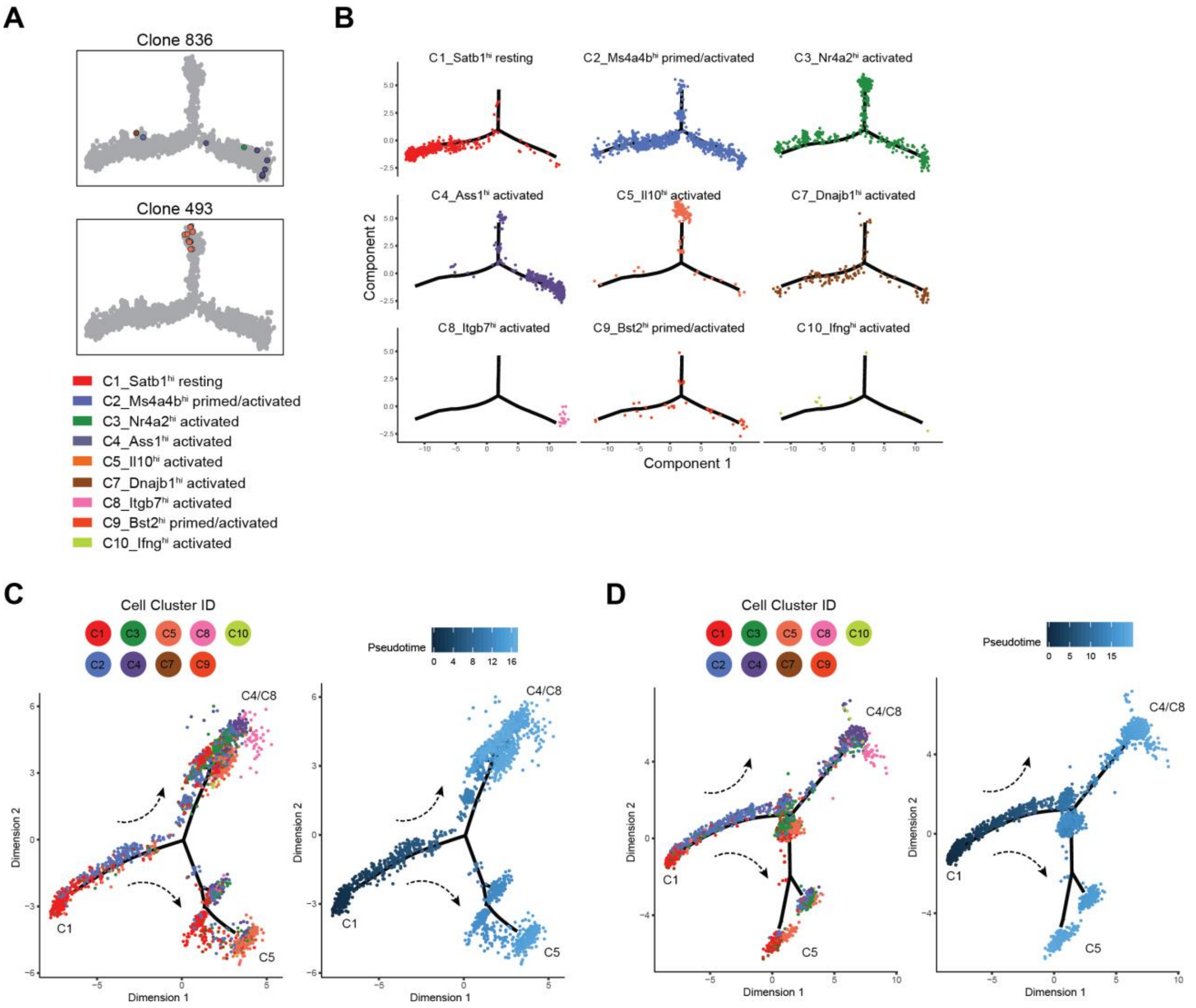


**Fig. S14. Trajectory analyses show consistent Treg differentiation patterns across replicates and gene expression dynamics along Treg differentiation.**

**(A)** Two representative clonal families involving Tnfrsf9$^+$Il1rl1$^+$ Tregs (Clonal family 836) and Il10-producing Tregs (Clonal family 493) are plotted along the trajectory manifold. Individual cells are colored by cell state. **(B)** Individual Tregs with recovered TCRs are grouped by Treg cell state and plotted separately along the trajectory manifold. **(C)** Pseudotemporal ordering of Tregs from replicate 1 (*n*=12,840 cells) without recovered TCRs. Individual cells are colored by cell state (left) or by pseudotime (right). **(D)** Pseudotemporal ordering of Tregs from replicate 2 (*n*=11,822 cells). Individual cells are colored by cell state (left) or by pseudotime (right).
